# A generative AI framework for disease-specific lung microtissue bioengineering

**DOI:** 10.64898/2026.04.15.718723

**Authors:** Ella Bahry, Jeanine C. Pestoni, Kai Hirzel, Taras Savchyn, Diana Porras-Gonzalez, Vera Getmanchuk-Zaporoshchenko, Martin Gregor, Thomas M Conlon, Ali Önder Yildirim, Kyle Harrington, Deborah Schmidt, Gerald Burgstaller, Michael Heymann

**Author notes:** These authors contributed equally (Ella Bahry, Jeanine C. Pestoni, Kai Hirzel). Equally contributing (Deborah Schmidt, Gerald Burgstaller, Michael Heymann). co-corresponding authors and joint supervision: Deborah Schmidt, Gerald Burgstaller, and Michael Heymann.

## Abstract

Generative Lung Architecture Modeling (GLAM) is an integrated bioengineering framework that couples high-resolution three-dimensional tissue imaging with generative artificial intelligence to de novo design and 3D-bioprint anatomically detailed lung microtissue models. Native extracellular 3D matrix architectures of pulmonary parenchyma were extracted from healthy, fibrotic, and emphysematous in vivo mouse disease models and processed through a computational pipeline containing pre-trained image segmentation and 3D mesh generation. The resulting datasets were used to train a U-Net generative diffusion model with attention layers capable of synthesizing healthy and diseased lung tissue architectures. Microtissue cubes of about 200 - 300 µm edge length of native and synthetic datasets were fabricated through high-resolution two-photon stereolithography with gelatin-methacryloyl biomaterial ink and successfully seeded with cells, demonstrating biological compatibility. In closing the loop between biological imaging, generative modeling, and high-resolution biofabrication, this integrated framework establishes generative AI as a functional design layer for tissue engineering. The resulting lung microtissues retained architectural features of the native and original tissues, making them an application-ready platform for customizable and scalable fabrication of biological tissue surrogates for preclinical modeling, drug testing, and precision regenerative bioengineering.

## Introduction

Homeostatic imbalances in the pulmonary system can be life-threatening by undermining gas exchange, structural integrity and immune defense. Chronic lung diseases, such as chronic obstructive pulmonary disease (COPD), pulmonary fibrosis (PF), and asthma, continue to pose significant health burdens globally ^1–3^. Especially, COPD remains among the leading causes of morbidity and mortality worldwide, with its prevalence still increasing ^2^. COPD is characterized by progressive and irreversible airflow limitation caused by structural remodeling of the small airways, chronic inflammation, and destruction of alveolar tissue, leading to emphysema and loss of lung elasticity ^4^. In contrast, PF is marked by excessive deposition of extracellular matrix (ECM) and progressive scarring around the alveoli in the lung, resulting in reduced gas exchange and eventual respiratory failure ^5^. Lung transplantation remains the only curative option for patients suffering from end-stage pulmonary diseases ^6^. Yet, vital, life-saving organ transplantation therapy is fundamentally limited by a persistent global shortage of donor organs and by suboptimal long-term outcomes due to chronic graft rejections or lifelong immunosuppression therapy ^7–9^. The high complexity of these diseases, combined with the critical shortage of suitable donor tissues for transplantation thus underscores the medical need for innovative solutions in regenerative medicine and bioengineering.

Traditional approaches in modeling diseases often rely on simplistic *in vitro* systems with limitations in replicating the complex human tissue architecture and its (patho)physiological responses ^10–12^. Animal *in vivo* models, while useful for studying certain aspects of diseases, frequently fail in translation due to physiological differences between animals and humans, but there are also growing ethical concerns about using animals in research ^13–15^. Especially in lung research, organoids and *ex vivo* precision-cut lung slices (PCLS) from murine and human tissue have emerged as powerful three-dimensional (3D) models for studying disease mechanisms and testing therapeutic candidates ^16–18^. *Ex vivo* PCLS are thought to bridge the gap between preclinical research and translation by preserving native 3D lung architecture, ECM composition, cellular diversity and cell-cell communication routes, however their application is restricted by limited tissue availability, high donor-to-donor variability and short-term viability ^19^.

Tissue- and bioengineering approaches including advanced 3D bioprinting hold promise to generate functional, ECM-mimicking scaffolds that can be customized to recapitulate a healthy or diseased organ microenvironment, and once scaled into fully functional organs, may provide solutions for organ transplantation, personalized medicine and regenerative therapies ^20,21^. Moreover, significant progress has been made in fabricating 3D-printed scaffolds that closely mimic *in vivo* ECM architectures ^22–24^. In particular, a demand to reconstruct tissue architecture with high-resolution drove the adaptation of two-photon stereolithography for bioprinting ^25^. Protein-based bioinks that incorporate gelatin methacryloyl (GelMA) further improved the biological functionality of these 3D-printed scaffolds markedly by providing biochemical cues to promote cell adhesion, spreading, and viability ^24,26,27^. Also, two-photon stereolithography could realize 3D-ECM microscaffolds of lung parenchyma that achieve native tissue biomechanics with tunable Young’s moduli, support colonization by primary human lung cells, and hold strong potential for advanced tissue engineering applications ^24^.

Artificial intelligence (AI) and in particular deep learning approaches continue to transform biomedical research and engineering by facilitating the analysis of large datasets and the generation of realistic data outputs ^28^. Among the foundational architectures, the U-Net - originally developed for biomedical image segmentation ^29^ - has become a workhorse for dense prediction tasks across modalities, including pre-trained models for volumetric tissue segmentation such as PlantSeg ^30^. Combined with self-attention layers, U-Net architectures now also form the backbone of modern generative diffusion models ^31,32^, which excel at synthesizing images from complex original datasets and have shown considerable potential in drug discovery, medical imaging, and the generation of synthetic datasets to overcome limitations of small or biased training data ^28,33,34^. Notably, diffusion-based inpainting has been applied to remove artefacts from 2D brain tissue image data ^34^, demonstrating that such models can learn disease-specific tissue patterns and generate structurally plausible tissue *de novo*. Concomitantly, AI is also increasingly integrated into 3D bioprinting workflows to optimize scaffold design, bioink formulations, printing parameters and quality control to improve reproducibility and functional outcomes ^35,36^. Extending beyond such print-process optimization, generative approaches such as GANs were also explored for synthesizing scaffold-like material structures ^37^. However, the integration of generative models trained on native tissue imaging data with high-resolution biofabrication has not yet been harnessed to produce disease-specific, biologically functional tissue scaffolds.

Here, we present a multidisciplinary approach of Generative Lung Architecture Modeling (GLAM) of human lung disease *in vitro* by closing the loop between biological data acquisition, AI-driven computational design and tissue engineering (**Figure 1**). COPD and fibrotic lungs from murine *in vivo* disease models were explanted and processed into 300 µm thick PCLS to preserve native lung architecture and extracellular matrix composition *ex vivo*. After decellularization and staining for ECM proteins a total of sixty 3D datasets comprising healthy and diseased lung parenchyma were acquired through 3D confocal microscopy. These spatial image datasets served as input for AI-based segmentation, followed by training of *in silico* generative diffusion models that learn disease-specific tissue structures and synthesize realistic and novel digital 3D tissue meshes. These meshes were then used as volumetric templates for *de novo* 3D bioprinting of realistic lung scaffolds via two-photon stereolithography with gelatin-methacryloyl hydrogels to mimic native ECM. Biological compatibility is demonstrated by successful colonization of the bioengineered scaffolds with human cells, thereby combining *in vivo*, *ex vivo*, *in silico* and *de novo* disease modeling. This continuous tissue engineering pipeline affords the opportunity to generate functional lung microtissue with custom anatomy to study disease dynamics, tissue remodeling and therapeutic responses, bringing us closer to our goal to reduce animal experiments and to ultimately overcome the organ donor bottleneck in curing severe pulmonary diseases.

**Figure 1:**
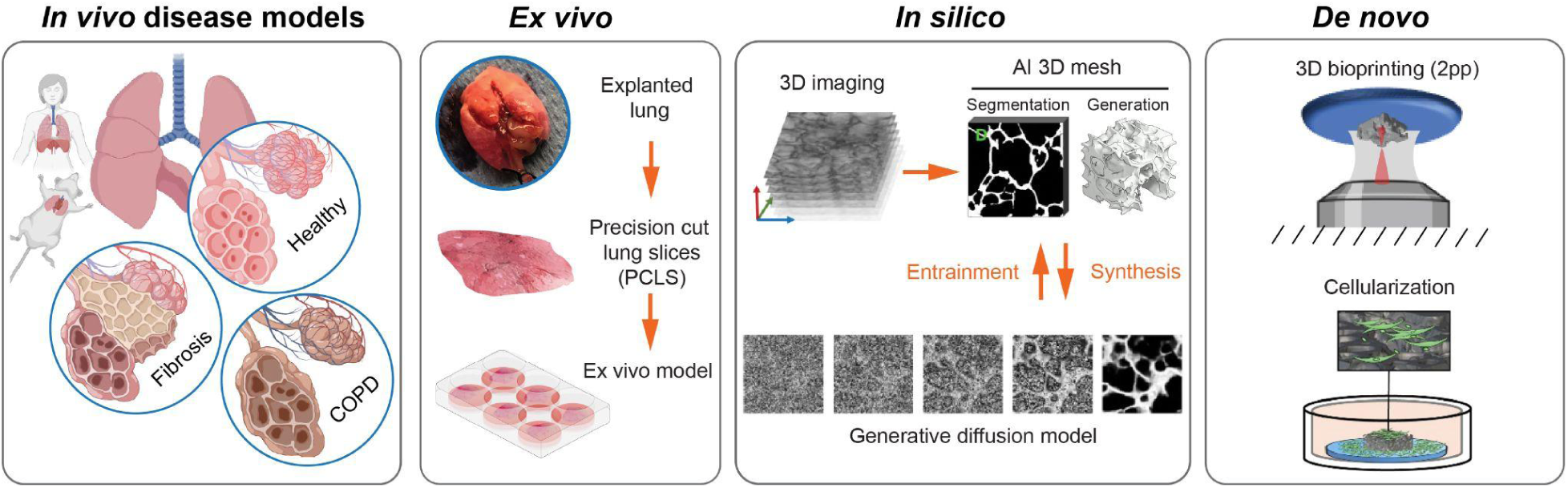
Integrated pipeline for lung disease modeling and AI-driven tissue engineering. Human chronic lung diseases like fibrosis and chronic obstructive pulmonary disease (COPD) can be modeled in animals (**in vivo**), thus enabling systemic investigations of disease pathogenesis and progression. Lungs from animals, but also from human donors, can be harvested and studied **(ex vivo**) as precision-cut lung slices (PCLS), an organotypic platform that preserves native 3D lung architecture, extracellular matrix composition, and cellular diversity. Digital reconstructions (**in silico**) of the lung’s microarchitecture is achieved by confocal high-resolution 3D imaging, which forms the basis for AI-based generative computational models trained on these spatial imaging datasets to produce anatomically realistic or disease-specific 3D meshes that subsequently can serve as templates for 3D bioprinting (**de novo**) of lung-tissue-scaffolds. Once printed, the scaffolds can be repopulated with human (stem) cells. Ultimately, this approach envisions the simulation of disease dynamics, tissue remodeling and therapeutic responses.

## Results and Discussion

### *In vivo* disease modelling and digital structural reconstruction

Investigating pathological features of chronic human lung diseases in animal *in vivo* models is well established ^38–40^. We replicated two widely used disease models: (1) Pulmonary fibrosis was induced by intratracheal instillation of bleomycin, which triggers idiopathic PF with epithelial injury, inflammation, and excessive extracellular matrix deposition; (2) Emphysema was modeled by oropharyngeal administration of elastase or by long-term exposure of cigarette smoke, both of which cause airspace enlargements due to tissue destruction characteristic of COPD; and (3) healthy control mice (**Figure 2A**). After surgical lung explantation, lungs were processed into and maintained as 300 µm thick *ex vivo* PCLS culture (**Figure 2B**).

**Figure 2:**
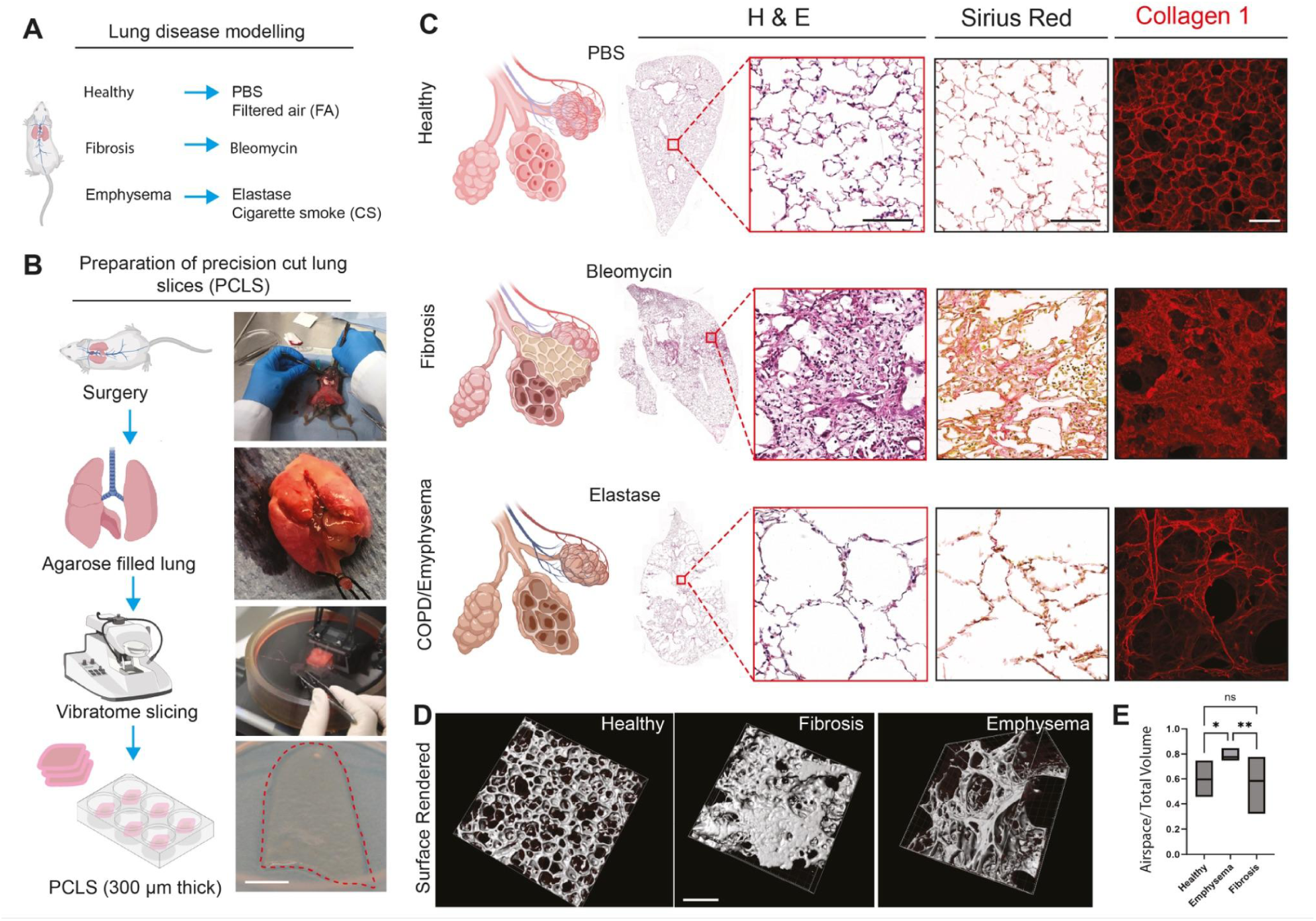
Disease modelling, generation of 3D precision-cut lung slices and histology. **(A)** Overview of in vivo mouse models for chronic lung diseases: healthy controls were treated with PBS or exposed to filtered air, fibrosis was induced by intratracheal bleomycin, and emphysema/COPD was induced via intratracheal elastase or cigarette smoke exposure. **(B)** Workflow for generating precision-cut lung slices (PCLS): after surgical lung explantation, lungs were filled with agarose to stabilize tissue structure. Lungs were then sectioned into 300 µm thick slices using a vibratome and transferred into multiwell plates. Scale bar = 600 µm. **(C)** Representative histological and immunofluorescent images of PCLS from healthy, fibrotic, and emphysematous lungs. Hematoxylin and eosin (H&E) stainings show parenchymal lung architecture. Sirius Red staining highlights collagen deposition, which is increased in fibrosis. Immunofluorescence staining with antibodies against Collagen-I (red) further indicates extracellular matrix deposition and remodeling across disease states. Scale bars = 100 μm. **(D)** Surface-rendered 3D reconstructions from 3D confocal imaging (z-stacks) reveal major structural alterations like tissue thickening in fibrosis and airspace enlargements in emphysema compared to healthy controls. Scale bar = 200 µm. **(E)** Quantification of the airspace-to-total-volume ratio confirms significantly increased airspace in emphysematous lungs (p*<0.05, p**<0.01) compared to healthy and fibrotic conditions. Statistical significance assessed by one-way ANOVA.

To obtain suitable organotypic lung parenchyma training data, we optimize the PCLS processing pipeline for artefact-free high-resolution imaging. For this, a baseline of physiological ultrastructure is obtained through histological staining with hematoxylin and eosin (**Figure 2C**). These confirmed intact alveolar parenchyma in healthy and control mice, dense fibrotic remodeling in bleomycin-treated mice, and enlarged, destroyed alveoli in emphysema models. In a second step, we access extracellular matrix architecture and lung tissue remodeling by selectively staining collagen fibers with Sirius red. Collagens are key structural components of the extracellular matrix and many isoforms are aberrantly overexpressed in lung fibrosis, while in COPD the extracellular matrix undergoes extensive degradation and remodeling ^4,41^. This analysis revealed increased collagen deposition in fibrotic lungs, while emphysematous tissue exhibited a reduced and more fragmented collagen network (**Figure 2C**). As the cigarette smoke model did not show consistent and reproducible structural changes compared to the elastase-administered mice, we hence focused our subsequent analyses on the latter as our emphysema model. PCLS were decellularized to improve their optical clarity for confocal imaging (**Supplementary Figure 1**). Removing cellular components reduces light scattering and hence fluorescence background. As a result, depth penetration during 3D imaging increased, affording an overall more accurate and deeper visualization of ECM features - especially for disease states like fibrosis with a dense ECM. Decellularized PCLS were shown to serve as a valuable model to study cell-matrix interactions in recellularization experiments, functional studies and high-resolution 3D-printing technologies ^24,42^. The extracellular matrix in decellularized PCLS was assessed by 3D confocal imaging after immunofluorescent staining for collagen type I. Disease-specific alterations in extracellular matrix organization were clearly reflected in these, similar to the previous Sirius red sections, suggesting that the physiological extracellular matrix architecture was preserved during decellularization (**Figure 2C**). Fibrotic lungs exhibited dense and continuous collagen I networks, consistent with excessive extracellular matrix deposition, where emphysematous tissues showed airspace enlargements and fragmented collagen I structures consistent with ECM degradation. Pathologically, these structural changes in the lung tissue are manifested as airflow restriction in lung fibrosis and air trapping in emphysema/COPD along loss of elastic recoil, both conditions severely impair gas exchange, increase progressive dyspnea, reduce exercise tolerance, and ultimately cause respiratory failure ^4,43,44^.

To further assess disease-specific alterations in lung architecture, we performed full-slice overview scans using confocal laser scanning microscopy on the collagen type I stained decellularized PCLS to identify representative regions of interest (**Supplementary Figure 2A**). These regions were subsequently imaged at higher magnifications as z-stacks ranging up to 300 µm in thickness to obtain 3D volume datasets. Consistent with previous results, these spatial datasets captured clear structural differences between healthy, fibrotic and emphysematous lungs and were further processed into 3D visualizations of extracellular matrix topology (**Figure 2D** and **Supplementary Figure 2B**). Quantification of their structural differences in 3D in airspace volumes (**Supplementary Figure 2C**) revealed a significantly higher airspace volume fraction in emphysematous lungs compared to both fibrotic and healthy lung tissue, reflecting the loss of structural integrity and alveolar wall destruction, which is characteristic of advanced COPD (**Figure 2E**). However, fibrotic samples displayed a higher variability in airspace volume fraction compared to healthy samples, likely due to disease-specific regional heterogeneity. Overall, our *in vivo* models reliably reproduced key structural and anatomical features of lung fibrosis and emphysema, which could then be captured in decellularized *ex vivo* precision-cut lung slices as well as subsequent 3D imaging thereof. A total of 60 3D datasets were acquired, which can hence serve as a reliable starting point for the AI-generation of anatomically accurate lung scaffolds (**Supplementary Table 1**).

### Generative diffusion model training and performance

Our computational pipeline processed high-resolution 3D confocal image stacks of murine lung tissue and trained a customized 3D generative diffusion model for synthesizing condition-specific lung architectures (healthy, emphysematous, and fibrotic) (**Figure 3**). Training progress was continuously monitored by tracking the validation loss and the foreground pixel ratio of generated volumes. As depicted in **Figure 4A**, the validation-loss moving average (smoothed with a 1000-epoch window) for all models exhibited an initial rapid decrease until approximately 5,000 epochs. Due to the limited number of training examples, the loss curves subsequently became noisy, characterized by fluctuations around a general downward trend that persisted until reaching their respective minima. Specifically, the emphysema model’s loss reached a minimum around 26,000-27,000 epochs, while the healthy and fibrotic models reached their minima around 17,000 and 18,000 epochs, respectively. These smoothed loss curves, particularly their overall trends and plateaus, were critical in determining the stopping criterion for training. Similarly, the foreground voxel ratio of the generated binary images matched closely with the baseline of the respective training sets (**Figure 4B**). For instance, the healthy model achieved a foreground ratio of 0.258 ± 0.063, which was comparable to the training set’s 0.247 ± 0.062. The emphysema model showed 0.181 ± 0.104 versus a training baseline of 0.185 ± 0.055, and the fibrotic model demonstrated 0.234 ± 0.101 versus a training baseline of 0.212 ± 0.052. The final models were selected based on the stability of these metrics, with the healthy model, for example, being chosen at epoch 17,000 (**Figure 4C** and **Supplementary Figure 3**).

**Figure 3:**
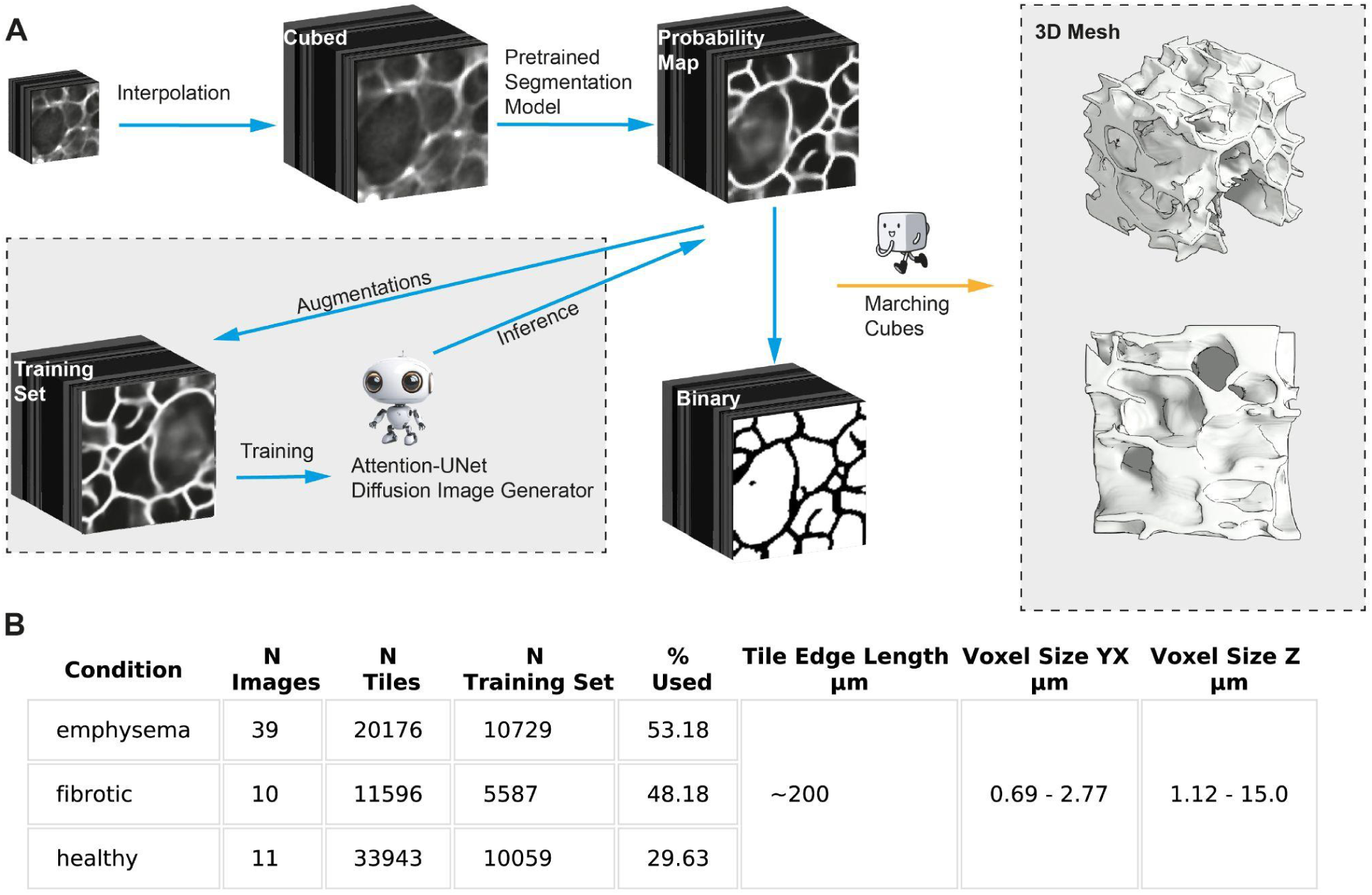
Digital workflow from raw data to printable meshes. **(A)** Original images are divided into 200×200×200 μm tiles and interpolated to isotropic resolution. Each lung tissue tile image is segmented using a pre-trained model, producing probability maps. These maps, along with their augmentations, form the training set for a diffusion model that generates synthetic image cubes per condition. Real and generated probability maps, together with their binarized versions, are converted into 3D virtual meshes for printing. **(B)** Table summarizing the number of original images per condition, resulting tiles, filtered tiles used for training, and tile dimensions.

**Figure 4:**
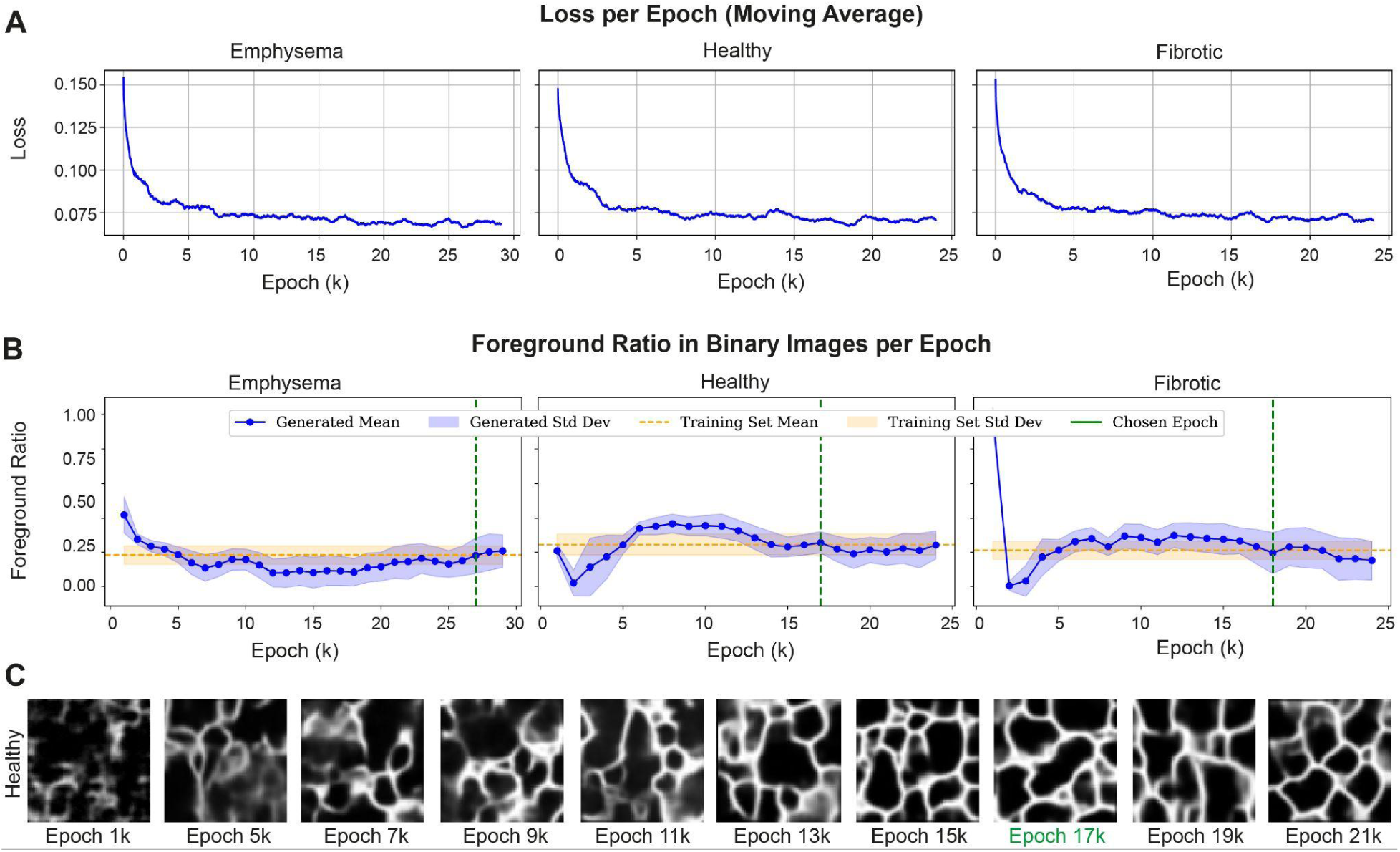
Training Process and Epoch Selection of Generative Models Across Disease Conditions. Statistical plots use 100 randomly sampled generated images; their metrics determine the stopping epoch. **(A)** The validation-loss moving average per condition is displayed across epochs, evaluated every 1,000 epochs. **(B)** The foreground voxel ratio per epoch for each condition shows generated binary images (mean ± SD) in comparison to the training set baseline. **(C)** An example 2D slice from a 3D tile generated by the healthy model is provided at selected epochs, with epoch 17,000 being the chosen final model.

### Validation of synthetic 3D architectures

To ensure the reliability and biological plausibility of both the input datasets and the generated 3D lung structures, a multi-tiered quality control framework was established, integrating quantitative, statistical, and structural assessments. The procedures were designed following recommendations emerging from recent benchmarking studies on validation of generative models in biomedical imaging and adapted to the specific context of diffusion-based 3D lung architecture synthesis ^45–48^. Quantitative comparison of the generated images against the original segmented data further confirmed the fidelity of the generative models. **Figure 5A** illustrates the mean pixel intensity comparison between generated probability maps and original probability maps, showing that generated images maintained similar intensity profiles across all conditions, although with slightly higher variance in generated outputs. Specifically, the mean pixel value for healthy generated images was 0.283 ± 0.066 compared to original 0.319 ± 0.013. For emphysema, generated was 0.228 ± 0.091 vs. original 0.301 ± 0.018, and for fibrosis, generated was 0.261 ± 0.096 vs. original 0.316 ± 0.011.

**Figure 5:**
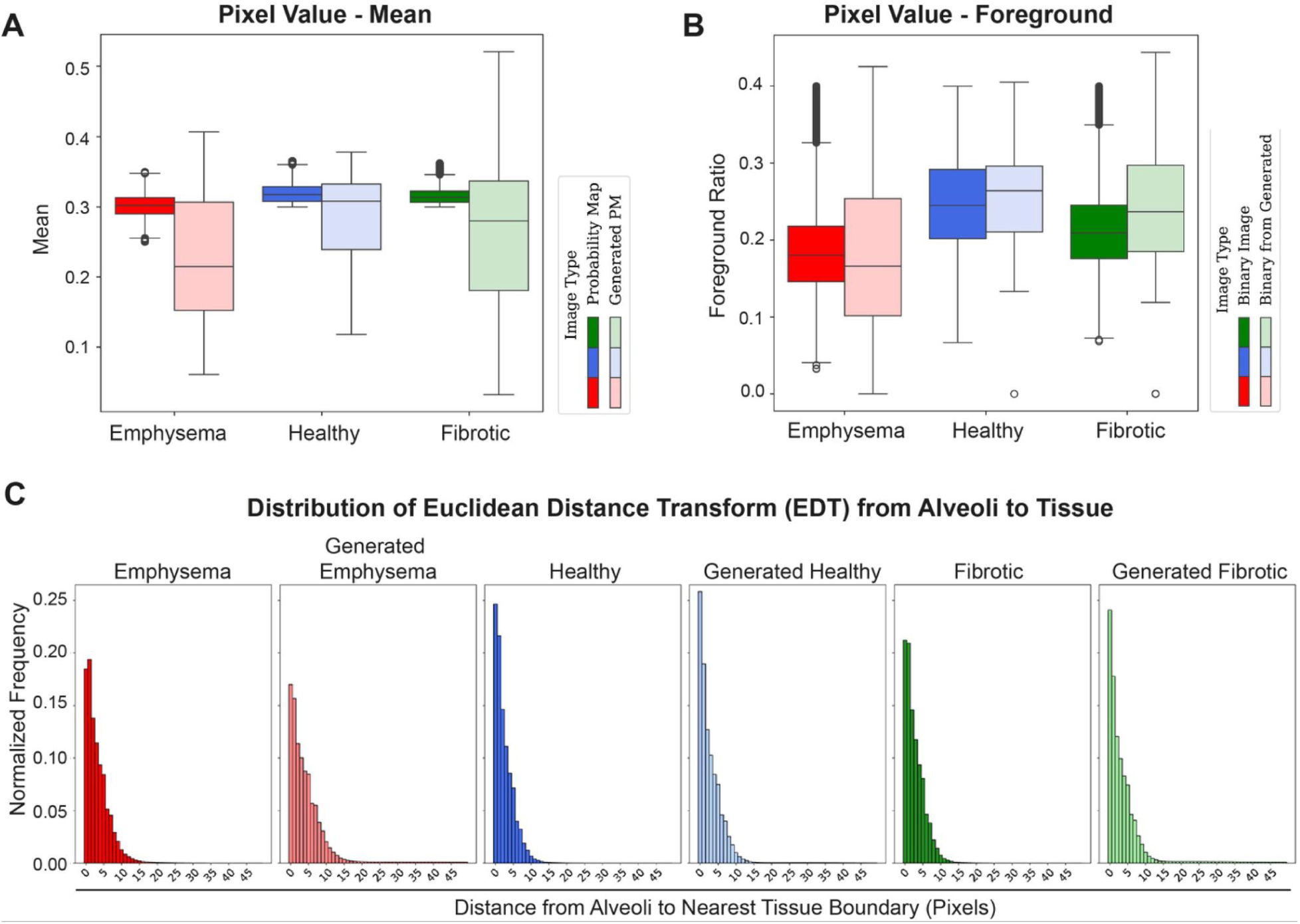
Statistical Comparison of Generative Model Performance Against Original Data. **(A)** Comparison of the mean pixel value between generated images and original data, presented per disease condition. **(B)** Ratio of foreground pixels in binary images (derived from the probability maps) for generated images versus original data, per disease condition. **(C)** Normalized pixel intensity histograms of the Euclidean distance transform from alveoli to tissue (using flipped binary images), comparing original data to generated images per disease condition. Statistical plots use 100 randomly sampled generated images. An example image and more information about Euclidean distance transform can be found in **Supplementary Figure 4**.

Further, the foreground pixel ratio in the binarized generated images, a critical metric for structural representation, closely mirrored that of the original data. As visually confirmed in **Figure 5B**, the generative models consistently replicated the volumetric proportion of lung structures across all conditions, demonstrating strong agreement between generated and original foreground ratios. To benchmark global realism and diversity of the synthetic datasets, we implemented several distributional and feature-space metrics recommended in recent diffusion-model evaluation frameworks. The three principal metrics - Fréchet Inception Distance (FID) ^45–47^, Generative Precision and Recall ^48^, and Multi-Scale Structural Similarity (MS-SSIM) ^47,48^ **-** quantify complementary aspects of model performance: global distribution alignment, feature-space fidelity and coverage, and image-level diversity.

FID measures the statistical distance between feature distributions in generated and original images by approximating them as multivariate Gaussians in a pretrained embedding space. We extracted 2048-dimensional features from 2D slices (every fifth slice) of each 3D volume using the penultimate layer of an **Inception-V3** network pretrained on **ImageNet**. Although this encoder originates from natural image datasets, it remains the de-facto standard for FID computation in a biomedical context and correlates robustly with expert quality ratings when applied consistently. Woodland et al. compared ImageNet- and domain-specific (RadImageNet) FIDs and found that ImageNet features yielded more stable, human-aligned scores, supporting our use of the canonical implementation ^45^. For our microscopy data, the per-class FIDs were **35.5 (emphysema)**, **45.6 (healthy)**, and **47.9 (fibrotic)**. In prior biomedical studies, histology and medical-imaging diffusion models typically report FIDs between **20–60** for realistic outputs ^46,47^. Our values therefore indicate that the generated lung architectures reproduce the statistical characteristics of microscopy data with high fidelity, particularly for the emphysematous condition.

Because FID alone cannot disentangle realism from diversity, we additionally computed **Precision** and **Recall** as proposed by Kynkäänniemi et al. ^48^. These metrics evaluate the overlap between the real and generated feature manifolds. **Precision** is the fraction of generated samples lying within the real-data manifold (fidelity) and **Recall** is the fraction of real samples represented within the generated manifold (coverage). Using k-nearest-neighbor radii (k = 3–10), we observed consistent trends of increasing overlap with larger k, as expected ^48^. At k = 10, the models reached **Precision ≈ 0.52 ± 0.04** and **Recall ≈ 0.42 ± 0.05**, with emphysema showing the most balanced fidelity and coverage. These values align with those reported for state-of-the-art diffusion models on histopathology datasets ^47,49^, where precision > 0.4 and recall > 0.3 generally denote convincing biological realism and diversity.

To quantify diversity directly in image space, we computed the **mean pairwise MS-SSIM** between randomly selected generated slices per condition. Lower values correspond to higher diversity. All classes exhibited low MS-SSIM values (**0.05–0.07** range), indicating that the generators do not produce redundant or over-smoothed outputs. These numbers are comparable to MS-SSIM < 0.1 reported for diverse histological generative models ^47,48^.

To further assess fine-scale structural integrity beyond pixel-wise metrics, we analyzed the Euclidean distance transform based distributions of alveolar distances, providing a robust measure of geometry and alveolar size variability. This analysis provides a robust measure of the size and distribution of alveolar spaces. As shown in **Figure 5C**, the distributions of distances for generated healthy lung structures closely resembled those of the original healthy tissue, with similar peak alignments and overall spread, suggesting that the model accurately replicated characteristic alveolar sizes. Similar trends were observed for the emphysematous and fibrotic generated tissues.

Further statistical analysis corroborated the fidelity of generative model performance to ground truth (**Supplementary Figures 4-6**). Interestingly, a subtle but consistent trend of higher frequency in lower Euclidean distance transform values was observed across all generated conditions compared to their original counterparts. This suggests that the generative models may introduce slightly more tissue, potentially manifesting as very small enclosed areas within the alveolar spaces. Subsequent analysis confirmed the presence of such small, isolated alveolar regions in the generated volumes (**Supplementary Figure 6**). However most of these were below 140 µm³ in size and thus discarded in downstream quantitative analyses. Taken together, the models accurately capture the characteristic alveolar size distribution of the original data, albeit minor topological artifacts were occasionally produced at the finest scales.

Finally, three-dimensional meshes for 3D printing were successfully created from both the original segmented probability maps and the synthetically generated volumes using the Marching Cubes algorithm. Meshes were consistently generated for all conditions, ensuring watertight and manifold representations through post-processing steps (e.g., removal of degenerate faces, fixing inverted normals, hole filling).

### 2-photon 3D bioprinting and scaffold colonization

Virtual tissue-derived and model-generated 3D mesh geometries were printed into gelatin methacryloyl bioink using 2-photon stereolithography (**Figure 6A**). Physiologically-scaled alveolar scaffolds were previously fabricated from a blueprint of a native murine lung tissue section using a GM10 bioink with LAP as a photoinitiator ^24^. Monitoring the 3D-printed lung tissue scaffold integrity by brightfield transmitted-light microscopy during fabrication and development revealed a high shape fidelity and that geometrical dimensions were maintained accurately throughout the process, without visible distortions along the x- and y-axis (**Figure 6B**). Accuracy along the vertical z-axis was assessed via multi-photon fluorescence microscopy, after staining scaffolds fluorescently with DAPI. Volumetric 3D microscopy image renderings confirmed the 340 x 340 x 140 µm large tissue-derived and the 200 x 200 x 200 µm large model-generated blocks to be consistent with their designed mesh geometries in 3D and across the full scaffold height without suffering from apparent deformations or collapsed sections (**Figure 6B**). Overall, the resulting 3D-printed hydrogel scaffolds were highly reminiscent of alveolar-airspace septal boundaries and surrounding ECM wall structures visually appeared to match the previously described lung parenchymal architectures of the different disease states. The observed wall thickness at times however exceeded the physiological thickness of about 2 µm ^50,51^. While the implemented two-photon process can in principle achieve sub-micron features, the highly curved alveolar surfaces require sufficient voxel overlap and hence dense scan-line placement to minimize unpolymerized gaps between them. To ensure this, we choose 0.5 µm for XY spacing and Z hatching direction. With this configuration, 3D mesh designs were hence well suited for two-photon polymerization of the GM10 bioink, as no additional support structures were needed for printing and fine gaps between individual laser scan paths were only noticeable at few select positions with particular high curvature (**Supplementary Figure 7**).

**Figure 6:**
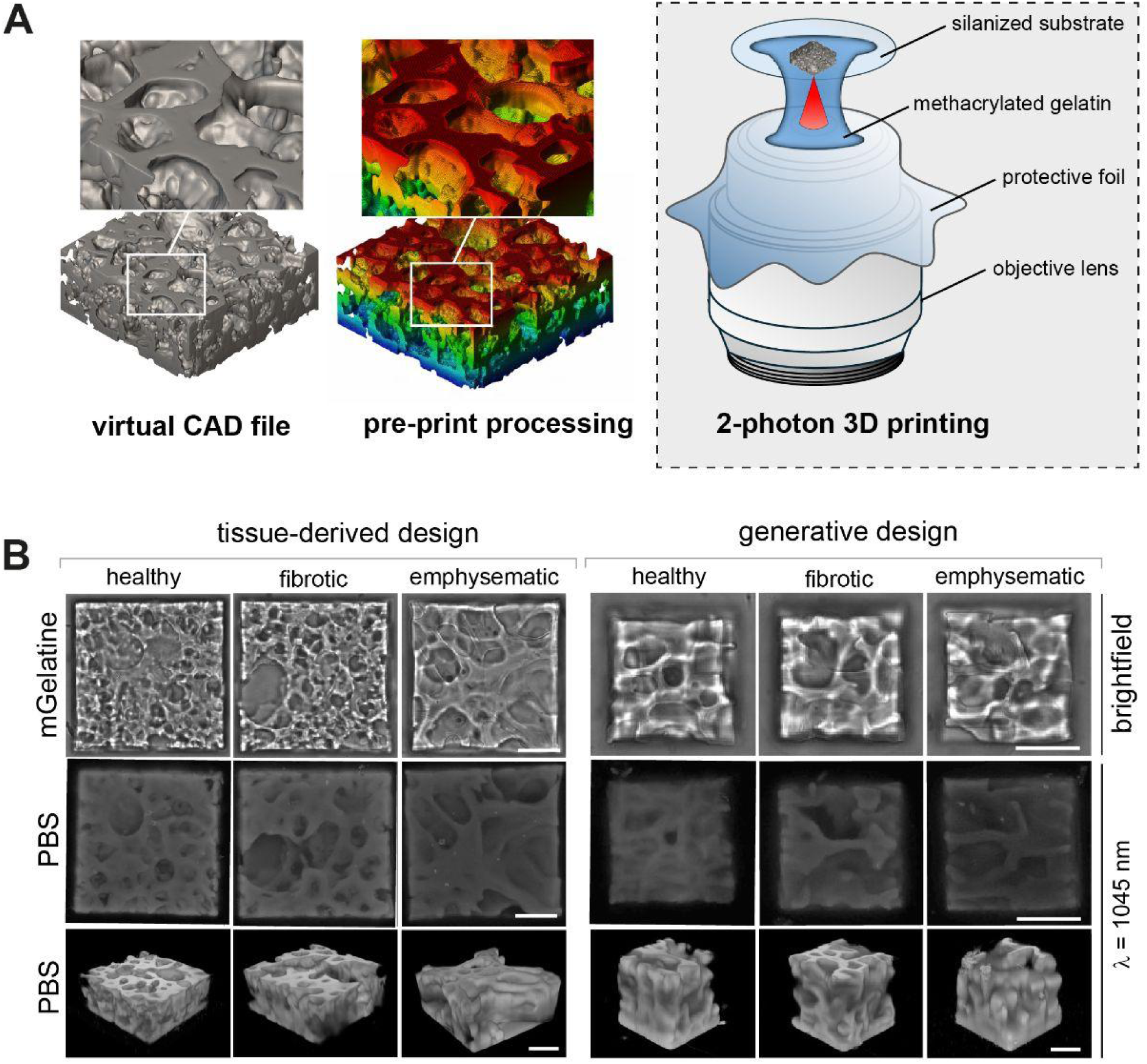
2-photon 3D bioprinting of tissue-derived and generative lung scaffold designs. **(A)** The virtual CAD files of the lung scaffold designs were processed pre-print for 500 nm hatching and slicing. Lung scaffolds were 3D bioprinted with 2-photon stereolithography using gelatin methacryloyl (GM10) in dip-in mode. **(B)** Brightfield images show top-down views of tissue-derived (left panel) and generative (right panel) lung scaffold designs immediately after printing in the GM10 print resin. 3D renderings of confocal z-stacks captured by two-photon imaging display a full 3D view of the 3D bioprinted scaffolds. Scale bars = 100 μm.

Finally, to assess the biological compatibility of the fabricated lung scaffolds and to demonstrate their ability to support cellular attachment and spreading, the scaffolds were transferred into standard cell culture dishes and colonized top-down with human MRC-5-GFP fibroblast cells (**Figure 7A**). Fibroblasts were selected as a biologically relevant cell type because they are principal producers and regulators of native ECM and its remodeling in the lung, contributing to tissue architecture during lung development, homeostasis, and repair ^42,52^. Importantly, fibroblasts establish a natural biologically conditioned matrix that provides structural, mechanical and biochemical instructive cues necessary for a subsequent integration and guidance of progenitor or stem cells. Here, scaffold-associated MRC-5-GFP fibroblasts were fixed after 24 hours of culture, stained with DAPI for cell nuclei and Phalloidin for intracellular cytoskeletal F-Actin, and imaged by 2-photon fluorescence microscopy (**Figure 7B**). Across all 3D-printed ECM scaffolds, fibroblasts attached and distributed uniformly along the alveolar septal- and vasculature wall-like regions. Partially, fibroblasts were even found lining luminal surfaces, implying that once adhesion and spreading of the cells is established, fibroblasts are capable of migrating along the scaffold’s surface. Surprisingly, compared with tissue-derived lung scaffolds, model-generated variants showed a slightly reduced distribution of attached cells across designs representing healthy, fibrotic and emphysematous states (**Figure 7B, right panel**). Of note, the 200 x 200 x 200 µm large model-generated scaffolds were substantially smaller than the 340 x 340 x 140 µm large tissue-derived designs, a difference that might contribute to the reduced number of adherent fibroblasts due to a reduction in the initial cell capture and adhesion during the seeding process (**Figure 7A**). Additionally, we previously identified and described small, closed alveolar regions in the generated volumetric designs (**Supplementary Figure 6**). These fine-scale topological artifacts are likely poorly accessible for cells and thus contribute to the observed reduction in cell attachment. Strikingly, scaffolds representing the emphysematous disease state showed the lowest level of cell attachment and cell density distribution, which might be the result of disease-related enlarged alveolar diameters or even alterations in the microtopological surface complexity, which together could limit early cell capture and adhesion during cell seeding (**Figure 7B**).

**Figure 7:**
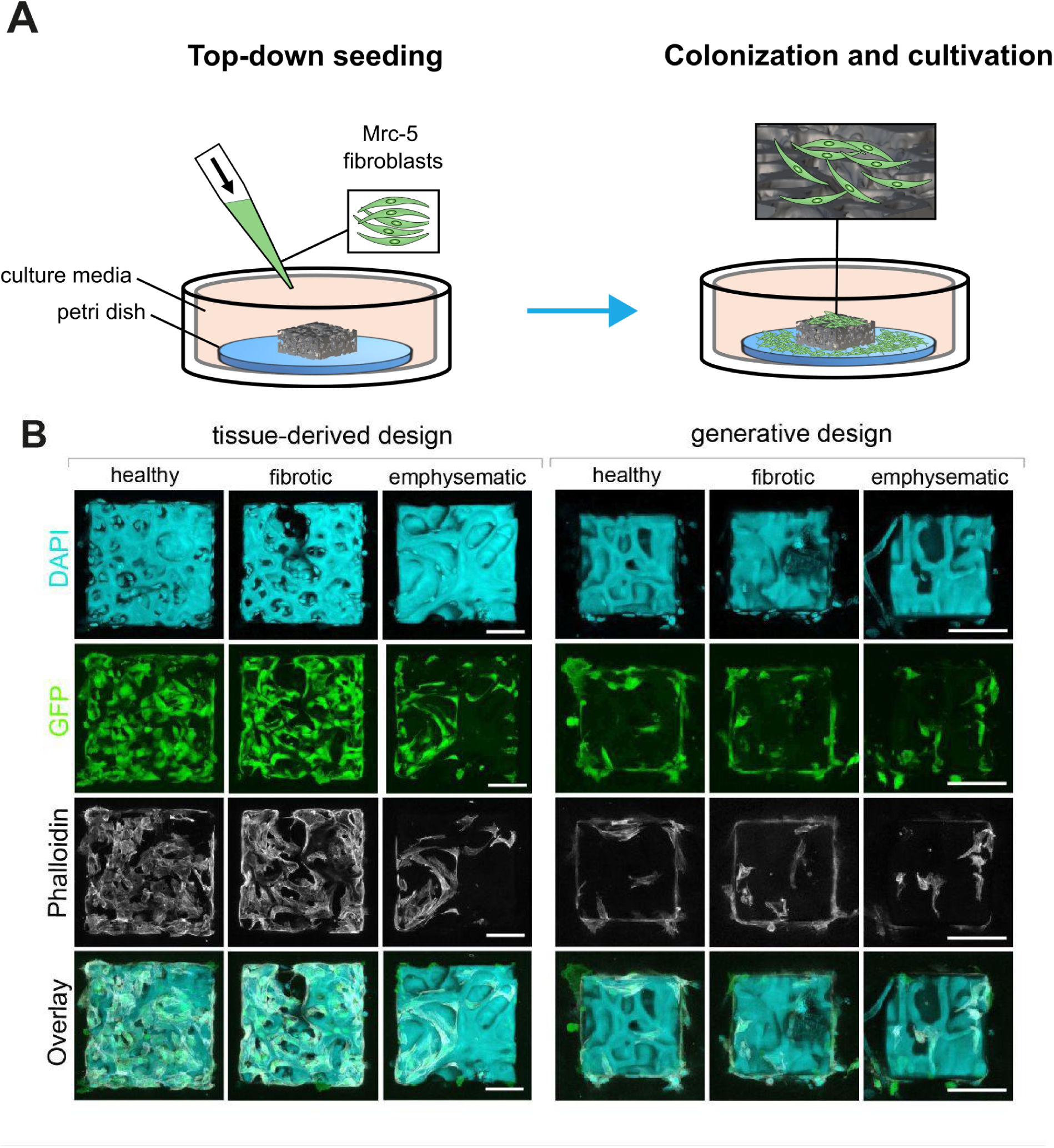
Tissue-derived and model-generated 3D lung meshes colonized with MRC-5-GFP fibroblast cells. **(A)** Workflow schematic for lung scaffold colonization. 3D bioprinted scaffolds were transferred to a petri dish with cell culture media and top-down seeded human with MRC-5-GFP fibroblast cells. **(B)** 24 hours post seeding MRC-5-GFP fibroblasts which were attached to the scaffolds were fixed and stained with DAPI (cell-nuclei, blue) and phalloidin (F-Actin, white). Notably, DAPI binds non-specifically to GelMA/GM10 of the printed scaffolds by electrostatic binding. Images show maximum-intensity projections of confocal z-stacks, revealing surface and luminal colonization of MRC-5-GFP cells (green). Scale bar = 100 μm.

The combination of high-resolution two-photon bioprinting and subsequent cellular colonization hence yielded biocompatible lung tissue surrogates that captured disease-specific geometries and lung-relevant microstructures, providing a structural basis for future integration of lung progenitor or stem cells. Intriguingly, it is reported that natural ECM can be highly instructive to cells, providing spatial, mechanical, and topological cues that regulate adhesion, migration, and cell fate ^42,53^. In line with this concept, our data indicate that the colonization efficiency of MRC-5-GFP fibroblasts in the bioprinted lung scaffolds is governed by the geometric accessibility, topology or scaffold surface exposure rather than by the intrinsic material properties of the GM10 bioink intended to mimic the native ECM architecture. These observations suggest that fibroblasts could function as a secondary biological layer of functional refinement, using the printed scaffold primarily as a geometric guide while depositing endogenous ECM that provides instructive cues for other functional lung-relevant cell types, which might be subsequently added.

## Conclusion

This study demonstrates an end-to-end pipeline that combines AI-driven structural synthesis with precision bioprinting for creating anatomically realistic microtissue models. In combining multidisciplinary state-of-the-art technologies from high-resolution 3D confocal imaging, AI diffusion-based generative modeling, and two-photon stereolithography, it was possible to recapitulate healthy, fibrotic and emphysematous matrix scaffolds colonized with human cells. The resulting 3D workflow integration of GLAM from 3D image to *de novo* 3D microtissue holds promise for pre-clinical drug testing, mechanistic studies, *in silico* simulations, digital inpainting by filling missing structural information, and above all, regenerative transplantation strategies across diverse tissues beyond the lung. These results establish generative AI as a viable computational design layer bridging biological imaging and tissue biofabrication. However, several limitations remain to be addressed.

A defining physiological function of the lung at the alveolar level is gas exchange across the thin air-blood barrier formed by alveolar epithelial cells, the intervening extracellular matrix, and a closely associated capillary blood network ^51^. Supporting perfusion and functional gas exchange in generated tissue models will hence be paramount in the future. Since bioprinted tissue scaffolds can capture lung-relevant architectures at the microscale of the finest vascular and capillary structures, their corresponding anatomy should be included in the training data. As our training data represents only limited regions within the lung parenchyma, larger-scale anatomical diversity such as airway branching patterns are consequently limited. Neuronal network-based image processing could recently skeletonize the fractal bronchial tree in embryogenesis ^54^. Cell type specific stains in turn could help to unambiguously discriminate vascular and capillary structures from alveolar septal ECM ^17^, but also high-resolution X-ray imaging ^55^ or volume electron microscopy ^56^ may all contribute adequate experimental ground truth data to ultimately entrain and synthesize fully functional lung tissues ^57^.

Despite their widespread use, optical imaging modalities such as confocal microscopy and multiphoton imaging, as used here, are particularly sensitive to signal loss from scattering in deeper sample regions. This effect was especially pronounced in dense fibrotic tissue, where a significant amount of acquired imaging data proved to be of insufficient quality and was excluded from model training. In the future, real-time image quality control during acquisition ^58^ will be invaluable for optimizing the allocation of acquisition time across samples and imaging settings. While anisotropic voxel sizes across our dataset were handled by cubic interpolation during preprocessing, self-supervised Z-slice augmentation ^59^ offers a more principled future alternative.

Further, tissue maturation reconciles biophysical forces with biochemical signaling. A complementary 3D mechanical tissue characterization via fluorescence microscopy ^60^ or acoustic manipulation and force microscopy ^61^ could further elucidate how mechanochemical feedback controls disease progression and tissue morphogenesis ^62^. For this, the GLAM pipeline may be coupled with geometric deep learning ^63^, or detailed biomechanical finite element tissue modeling such as SimuCell3D ^64^ or MorphoMechanX ^65^.

Generative methods remain computationally expensive, which limits the synthesized model volume to below a cubic millimeter. Therefore, improving computational efficiencies will help to realize larger tissue constructs. Recent advances in fast feed-forward 3D reconstruction suggest that 3D synthesis time can be substantially reduced while maintaining output quality ^66^.

Additive manufacturing remains amongst the most expensive fabrication technologies, especially in bioengineering. The printed scaffolds presented here are also restricted in size, thus high-resolution 3D bioprinting technologies have to be developed that support the fabrication of larger scaffolds. This is due to the sequential line-wise printing of the two-photon process used here ^24^. Volumetric printing alleviates the need for sequential fabrication and hence enables unprecedented fabrication speeds of a few seconds, but not yet similar resolution ^67^. Above all, correct placement of functional cells and matrix embedding within a tissue remains a major challenge in bioengineering. Scaffold colonization in this study was limited to fibroblasts and did not account for the complex multicellular architecture of organs. Aside from instrumentation innovation, responsive bio-inks could close this resolution gap by shrinking into shape post fabrication ^68^.

Finally, safety must be of paramount concern when deploying generative AI in a medical context. Clinical applications of GLAM - including diagnostics, drug testing, or ultimately therapeutic tissue fabrication - will require rigorous validation before translation. Generative diffusion models are known to produce hallucinations: structurally plausible but biologically incorrect outputs that may not be immediately apparent yet could lead to flawed downstream results ^69^. The small enclosed alveolar regions we observed in generated volumes are a direct example of such artifacts in our own pipeline. Establishing robust, multi-tiered validation frameworks that assess not only distributional fidelity but also task-level biological plausibility ^70^ will therefore be essential for any clinical use of AI-driven tissue fabrication.

Overcoming these limitations could facilitate the deployment of clinically relevant tissue and organ surrogates. A particularly exciting prospect would integrate generative tissue modeling into an endoscopic bioprinter to facilitate tissue regeneration directly inside the patient, bypassing organ transplantation entirely, as a range of intravital printing techniques are actively developed ^71–73^.

## Methods

### Animal Experiments

Eight- to twelve-week-old C57BL/6N male/female mice were used in all experiments. Mice were housed under specific pathogen-free conditions and maintained at a constant temperature (20 - 24 °C) and humidity (45 - 65 %) with a 12 hour light cycle, and were allowed food and water ad libitum. Mice were euthanized by terminal exsanguination following anesthesia. The following disease models, including various control conditions were used:

**(1) Bleomycin-induced pulmonary fibrosis mouse model:** After anesthetization with 50 µl of Zoletil/Rometar by intramuscular injection, mice were given a single dose of 3U/kg Bleomycin (Sigma-Aldrich) by intratracheal instillation of 1.2 U/ml in phosphate buffered saline (PBS), e.g. 50 µl/20 g mouse. Control mice were instilled with 50 µl of PBS/20 g mouse body weight. On day 14 after bleomycin administration mice were sacrificed and lung tissues were harvested for subsequent experiments.
**(2) Cigarette smoke-induced emphysema model:** Cigarette smoke was generated from 3R4F Research Cigarettes (Tobacco Research Institute, University of Kentucky, Lexington, KY), with the filters removed. Mice were whole-body exposed to active 100% mainstream cigarette smoke of 500 mg/m^3^ total particulate matter for 50 min twice per day for 3 days or 4 months, in a manner mimicking natural human smoking habits as previously described ^38,39^. The total particulate matter level was monitored via gravimetric analysis of quartz fiber filters prior to and after sampling air from the exposure chamber and measuring the total air volume. Filtered air exposed mice were used as controls. Mice were analyzed the day after the final smoking exposure.
**(3) Elastase-induced emphysema mouse model:** Emphysema was induced in mice by oropharyngeal application of a single dose of porcine pancreatic elastase (PPE, 40 U/kg body weight in 80 µl volume, Sigma-Aldrich) as previously described ^40^. Control mice received 80 µl of sterile PBS. Mice were analyzed 28 days after instillation.

All experiments were performed in accordance with the guidelines of the Ethics Committee of the Helmholtz-Center Munich and approved by the Regierungspräsidium Oberbayern, Germany or in accordance with an animal protocol approved by the Animal Care Committee of The Institute of Molecular Genetics and according to the EU Directive 2010/63/EU for animal experiments.

### Generation of murine precision-cut lung slices (PCLS)

Healthy, fibrotic, and emphysematous mice were anesthetized using a mixture of ketamine (Bela-Pharma) and xylazinhydrochloride (Cp-Pharma). Following intubation and diaphragm dissection, the lungs were perfused via the heart with sterile PBS. Using a syringe pump, warm low-melting agarose 3% (w/v) (Sigma) dissolved in sterile DMEM-F-12 (GIBCO) supplemented with penicillin-streptomycin and amphotericin B (both Sigma), was infused into the lungs. The trachea was closed with a thread to keep the agarose inside the lung. Subsequently, the lung was excised, placed into a cultivation medium, and cooled on ice for 10 min to allow agarose gelling. The lobes were separated and cut with a vibratome (Hyrax V55, Zeiss) to a thickness of 300 µm. Obtained murine PCLS were immediately processed for decellularization, histology or immunofluorescence.

### Decellularization of murine PCLS

Decellularization of murine PCLS proceeded as previously reported ^42^. Briefly, 300 µm thick PCLS were washed three times for 5 min in sterile deionized water, then incubated in 50 ml deionized water in Falcon tubes for 16 hours at 4°C on a tube roller. After additional rinsing in deionized water, the PCLS were incubated in a 50-ml, 0.1% (w/v) SDS solution for 4 hours at room temperature. After washing twice in deionized water for 10 min each, the PCLS were incubated in 1 M NaCl for 16 h at 4°C on a tube roller. After washing twice in deionized water for 10 min each, the PCLS were incubated in 7.5 ml PBS together with 5 mM MgCl_2_ and 30 µg/ml DNAse for 3 hours at 37°C. Finally, the PCLS were washed three times in deionized water for 10 min each and stored in 24-well plates (TPP Techno Plastic Products) containing PBS that was supplemented with penicillin-streptomycin (Sigma).

### Immunohistochemistry of decellularized PCLS, confocal fluorescence microscopy, and 3D image analysis

Decellularized PCLS were washed twice in PBS containing 138 mM NaCl, 26 mM KCl, 84 mM Na_2_HPO_4_, and 14 mM KH_2_PO_4_, pH 7.4. Primary Collagen I (Rockland) antibody was diluted in 1% (w/v) bovine serum albumin (BSA; Sigma) in PBS and then incubated with decellularized mPCLS for 16 hours at 4°C, before washing three times with PBS for 5 min each. Secondary antibody stainings with Alexa-fluor anti rabbit 568 (Thermo Fisher Scientific) followed the same protocol. For imaging, stained decellularized PCLS were transferred into a glass-bottomed, 35-mm CellView cell culture dish (Greiner BioOne). A drop of PBS was added on top of the decellularized PCLS, a piece of moistened paper tissue was placed around the periphery of the dish, and the lid was tightly sealed with Parafilm to prevent dehydration. Lung slice images were acquired as z-stacks, using an inverted confocal LSM 710 (Zeiss) with 10x/0.45 NA, 20×/0.8 NA and 40× /0.95 NA Plan-Apochromat, operated with ZEN2009 software (all Zeiss). Imaris 9.5 software (Bitplane) was used to render volume and surface views of the obtained confocal fluorescent z-stacks, and to quantify 3D volumes of the extracellular matrix and airspace via its Measurement Pro tool.

### Formalin-fixed paraffin-embedded (FFPE) sections and histochemistry of mPCLS

mPCLS were fixed in 4% paraformaldehyde (pH 7.0) overnight at 4 °C. Fixed slices were placed in embedding cassettes and processed using a Microm STP 420D tissue processor (Thermo Fisher Scientific) with two 60-min incubations in 4% formalin, followed by graded ethanol dehydration (50%, 70%, 96%, and 100%; each for 60 min, with 96% and 100% repeated twice) and paraffin infiltration (one 30-min and three 45-min cycles). Samples were then embedded in paraffin Type 3 (Thermo Fisher Scientific) using the EC 350 modular embedding system (Thermo Fisher Scientific), and the resulting blocks were stored at 4 °C. FFPE sections of 3 µm thickness were cut on a Hyrax M55 microtome (Zeiss, Germany), mounted on glass slides, dried overnight at 40 °C, and stored at 4 °C. Deparaffinization and rehydration were performed sequentially in xylene (2 × 5 min), 100% ethanol (2 × 3 min), 90% ethanol (3 min), 80% ethanol (3 min), 70% ethanol (3 min), and Milli-Q water (5 min). Next, these sections were stained with Sirius red (13422; Morphisto) and Hematoxylin (T865; Roth) according to the manufacturer’s instructions or with Hematoxylin (T865; Roth) and Eosin Y solution (0.5%; X883; Roth). Imaging of total FFPE-sections from multiple slides was accomplished by using an Axioscan 7 equipped with a 20x/0.8 NA Plan-Apochromat objective, operated with ZEN2012 software (all Zeiss). The final images were analyzed, stitched, cropped and individually adjusted for contrast and brightness by using ZEN2012 software (Zeiss).

### Image Analysis

Three-dimensional image stacks were processed in a multi-step image analysis pipeline designed for volumetric lung tissue reconstruction. This pipeline, glampipe, included image preprocessing, semantic segmentation, and subsequent 3D mesh generation. The images were also used for the training and application of a 3D image generation diffusion model. The full digital workflow can be found in **Figure 3A**.

### Image preprocessing

Original image stacks exhibited varying anisotropic voxel sizes, ranging approximately 2.3 to 15 µm in depth (Z), and 0.68 to 2.76 µm in lateral (XY) directions, were first processed to ensure consistent dimensions for downstream analysis (**Figure 3B, Supplementary Table 1**). A summary of all image processing parameters can be found in **Supplementary Table 2**.

1. **Tiling:** Image stacks were divided into 3D cubes with an edge length of approximately 200 µm. The voxel dimensions of these lung cube tiles varied depending on the original image’s voxel size. Tiles containing insufficient signal or deemed empty were excluded from further analysis.
2. **Voxel resampling and interpolation:** To achieve pseudo-isotropic voxels and a consistent resolution, images were resampled using cubic interpolation (scipy.ndimage.zoom). As the number of images was limited, we aimed not to omit data with low Z-resolution. Also, the XY- and Z-resolutions diverged by up to fivefold, which we consider a technical limitation of the available microscopy rather than a biological ground truth inherent to the underlying lung tissue architecture. Hence, together with image augmentations of 90° rotations across all axes, interpolation allowed us to train a rotation- and axis-agnostic image generator diffusion model. For down-sampling operations, a Gaussian filter was applied as an anti-aliasing measure. Interpolation factors were chosen for each image according to original voxel dimensions, with factors such as (3.5,2.0,2.0), (0.5,0.5,0.5), (6.5,2.0,2.0), (4.5,2.0,2.0), (5.0,2.0,2.0) being typical examples. After interpolation, all tiles were resized to a final shape of 92×92×92 voxels with ∼200 µm per edge.
3. **Contrast enhancement:** Prior to segmentation, image contrast was uniformly enhanced across the 3D stacks via histogram equalization. For this, the cumulative distribution function (CDF) of the entire 3D image was computed and normalized pixel intensities were mapped to ensure optimal signal range utilization.
4. **Semantic segmentation:** Segmentation was performed to identify structures of interest via the pre-trained deep learning model PlantSeg lightsheet_3D_unet_root_ds2x from SNZLab/PlantSeg ^30^. This model was chosen from several tested pre-trained 3D segmentation networks, as it produced the most faithful segmentations of the complex lung branching architectures in our datasets. Both lung tissue and plant roots are organized into fractal branches. In contrast, some otherwise promising models generated artifacts such as a “pancake defect” in the Z-direction (**Supplementary Figure 8A**), likely due to their inference on smaller patches. To choose a segmentation model, a small image set was annotated by two different individuals to obtain manual ground truth annotations, which were compared against the predictions of each model (**Supplementary Figure 8B**). The chosen PlantSeg model is a variant of a 3D U-Net trained on light-sheet images of *Arabidopsis* lateral roots at 1/2 resolution, with a voxel size of 0.25×0.325×0.325 µm (ZYX), and was trained using a BCEDiceLoss. It generated voxel-wise probability maps, which indicate the likelihood of each voxel belonging to the target structure. These probability maps served a dual purpose: 1) directly as input for subsequent mesh generation, and 2) after specific image augmentation, as a training set for the generative diffusion model.
5. **Binarization:** The generated probability maps were subsequently binarized using Otsu adaptive thresholding ^74^. Otsu’s method automatically determines an optimal global threshold to separate foreground from background based on the image histogram. The resulting binary masks were used together with the probability maps for 3D mesh generation.

### Diffusion model for image generation

Following segmentation, 3D image generation was performed using a customized three-dimensional (3D) diffusion model, which was trained on the segmented probability maps. This model was adapted from the lucidrains/video-diffusion-pytorch framework ^75,76^ by replacing the original 2D + time attention layers with a single 3D spatial attention layer - an essential modification for processing 3D volumetric data (**Supplementary Figure 9**). To accommodate grayscale inputs, the initial input layer was adapted to accept a single channel instead of the standard three RGB channels. The model architecture, Unet3D, was initialized with a base dimension dim = 64 and dim_mults = (1, 2, 4, 8), to configure a multi-resolution U-Net-like structure. The diffusion process was implemented as a GaussianDiffusion model with timesteps = 1’000. Due to memory constraints, the input image was sized to 80×80×80 voxels, downscaled from the 92×92×92-voxel interpolated tiles used in previous steps. An L1 loss was used to optimize the diffusion model during training.

#### Model training

A separate diffusion model was trained for each experimental condition (emphysema, fibrotic, and healthy). Each model was trained on a corresponding dataset of augmented 3D probability maps. The training data was generated with 100 iterations of augmentation using torchio to create a diverse dataset ^77^. The augmentation pipeline included intensity rescaling to a [0, 1] range (tio.RescaleIntensity(out_min_max=(0, 1))), random elastic deformations parameterized by num_control_points=(5, 5, 5), max_displacement=(4, 4, 4), and locked_borders=1. Additionally, random affine transformations (tio.RandomAffine) were applied, with fixed scales (scales=(1, 1)), rotations chosen from 0,90,180, or 270 degrees (in 90-degree multiples independently for X, Y, and Z axes) while preserving isotropic voxel spacing (isotropic=True) and rotating around the image center (center=’image’). Finally, random flips along any of the three axes (tio.RandomFlip(axes=(0, 1, 2))) were also incorporated. This comprehensive data augmentation strategy, particularly the random 3D rotations, was developed to mitigate residual resolution anisotropy to allow the diffusion model to generate rotation-agnostic volumes.

Training was conducted using the Adam optimizer with a learning rate of 1×10−4. A train_batch_size = 1 was used with a gradient_accumulate_every = 2 to effectively increase the batch size. Training proceeded for train_num_steps = 30’000 steps. An exponential moving average (EMA) of model parameters was maintained with ema_decay = 0.995. Mixed precision training (amp = True) was utilized to accelerate the training process and reduce memory consumption. Intermediate samples and model checkpoints were saved every save_and_sample_every = 1’000 steps. The stopping criterion for training was determined by monitoring the validation loss and the ratio of foreground pixels in the generated volumes, matching these statistics to those of the training set. To mitigate noise and discern trends, a moving average with a window size of 1’000 epochs was applied to the raw validation loss data. A summary of all image processing parameters can be found in **Supplementary Table 3**.

### 3D mesh creation for bioprinting

Three-dimensional meshes representing the segmented structures were created from probability maps and their binary counterparts of both the original segmented tissue, as well as from the diffusion model generated volumes. This process ensured consistent mesh representation across both experimental and synthetically generated data. A summary of all mesh generation parameters can be found in **Supplementary Table 4**.

1. **Marching cubes algorithm:** Meshes were generated using the Marching Cubes algorithm (skimage.measure.marching_cubes) ^78,79^. For this process, probability map images and their corresponding binary masks were padded by 5 pixels to avoid boundary artifacts. Binary dilation (scipy.ndimage.binary_dilation, 3 iterations) ensured robust mesh generation, particularly for small or thin structures. An optimal threshold (isolevel) was derived from the same Otsu threshold applied during binarization.
2. **Mesh post-processing:** Post-processing using the trimesh library ^80^ included degenerate faces removal (mesh.unique_faces(), mesh.nondegenerate_faces()), general mesh processing (mesh.process()), fixing inverted normals (mesh.fix_normals()), and filling any holes (mesh.fill_holes()). These steps ensured watertight, manifold meshes - free of gaps or self-intersections - suitable for bioprinting.
3. **Hardware:** Model training was performed on a single NVIDIA A40 GPU with 48 GB of VRAM.

### High-resolution 2-photon 3D bioprinting using a custom dip-in setup

High-resolution 2-photon 3D printing was performed on a Photonic Professional GT2 equipped with a 25x 0.8 NA objective (all Nanoscribe GmbH). Prior to the print process, virtual 3D mesh files were pre-processed for hatching and slicing (both 500 nm) using Nanoscribe ‘DeScribe’ software. Virtual 3D meshes were 3D-printed with a laser scan speed of 15’000 µm s^-1^, laser center wavelength at 780 nm and an average output power of 60 mW. To prevent bioink dehydration during 3D printing, a custom water-filled adapter ring was mounted around the objective lens to maintain a locally humidified environment (**Supplementary Figure 10**). Tissue-derived and model-generated 3D meshes were 3D-printed on round glass substrates (neoLab Migge GmbH, 20 mm diameter, 170 µm thickness). To improve attachment of the 3D-printed scaffolds to the substrate surface, the glass substrates were pre-treated for 15 min with a 1:200 dilution of 3-(trimethoxysilyl) propyl methacrylate silane in 100% ethanol, then rinsed with 100 % ethanol and subsequently dried under a stream of nitrogen. The objective lens was prepared with transparent foil for wrapped writing mode printing as previously described ^81^. A drop of the hydrogel precursor solution of approximately 20 µl, 25% w/v GM10, 2% LAP in H_2_O was then each applied to the center of the glass substrate surface and onto the top of the print objective lens. Both drops connected upon final assembly of the glass substrate within the print device and formed a single print solution volume residing on top of the objective lens during the print process. Resulting 3D prints were extensively washed with H_2_O to remove uncured hydrogel precursor solution.

### Cell cultivation and lung tissue scaffold colonisation

Embryonal lung tissue fibroblast cells MRC-5 stably transfected with GFP were cultivated in RPMI1640 growth media supplemented with 10 % FCS and 1 % penicillin/streptomycin in a humidified cell culture incubator at 37 °C and 5 % CO_2_. Colonization of MRC-5-GFP cells on 3D-printed lung tissue structures was performed by top-down adding a cell suspension of 500’000 cells mL^-1^ in RPMI 1640 growth medium inside a 6-well plate. After 24 hours post cell seeding, adhered MRC-5-GFP cells were fixed with 4 % PFA for 15 min at RT, permeabilized with 0.1 % Triton-X100 for 5 min, and subsequently stained for immunofluorescence for 60 min at RT using Phalloidin-AF546 and DAPI.

### 2-photon microscopy of colonized lung tissue scaffolds

High-resolution and in-depth 2-photon imaging was performed using the Leica Stellaris 8 microscope by upside-down, dip-in mode using a 25x 0.8 NA water immersion objective (LEICA). Z-stack confocal images were obtained by excitation light of 1045 nm for DAPI (3-photon absorption), 980 nm for GFP (2-photon absorption) and 1200 nm for Phalloidin-AF546 (2-photon absorption).

## Acknowledgements

The Burgstaller lab is grateful to Marisa Neumann for excellent technical assistance. We all thank Dagmar Kainmüller for reviewing and editing the initial funding proposal and Polychronis Stamakis for manual ground truth annotations. The authors gratefully acknowledge the Technology Platform “Cellular Analytics” of the Stuttgart Research Center Systems Biology for their support and assistance in this work. We thank Stephan Eisler and Melanie Noack for their technical assistance with multiphoton imaging.

## Data and code availability

The volumetric lung tissue reconstruction datasets are available on Zenodo at https://zenodo.org/records/17280675. The image analysis pipeline, glampipe, is publicly available on GitHub at https://github.com/ida-mdc/glampipe. The code for the 3D image generation diffusion model is available on GitHub at https://github.com/ida-mdc/diff3d. Notebooks generating the computational results and plots can be found at https://github.com/ida-mdc/GLAM-paper-plots.

## Author Contributions

**EB:** Developed the 3D image analysis pipeline, including virtual 3D mesh generation and the training of a novel generative diffusion model that she and **TS** adapted for 3D datasets. She also performed the quantitative validation of the results and wrote and critically revised the manuscript. **JP:** Performed animal experiments including lung isolation, *ex vivo* PCLS production, and subsequent cell and tissue culture work. Performed 3D confocal microscopy, data generation, image analysis, image processing and rendering. Contributions to the manuscript. **KaH:** Mesh processing for 3D printing, 2-photon 3D printing and development, cell colonization of scaffolds, 2-photon imaging to verify the 3D prints and the colonization. **DPG:** Contributions to the manuscript writing, figure assembly, design and visualization. **VG-Z, MG, TC, AÖY:** Performed animal experiments, interventions and provided resources. **KyH:** Conceptualization, funding acquisition, and advising on the computational pipeline. **DS:** Funding acquisition, supervision of the computational part, data analysis, project administration and coordination, contribution to the manuscript. **GB:** Conceptualization and study design, funding acquisition, supervision of the *ex vivo* experimental part, project administration and coordination, providing resources, data analysis and interpretation, manuscript drafting, writing, and critical revision. **MH:** Conceived and initiated the project. Conceptualization and study design, funding acquisition, supervision of the biofabrication part, project administration and coordination, providing resources, data analysis and interpretation, manuscript drafting, writing and critical revision. All authors read and corrected the final manuscript.

