## Supplementary Information for "A generative AI framework for disease-specific lung microtissue bioengineering"

* These authors contributed equally.

### Supplementary Tables

#### Supplementary Table 1: Original Images and Metadata

| File Name | Condition | N Tiles | Voxel Size YX µm | Voxel Size Z µm |
|---|---|---|---|---|
| 40x_Fibrotic_ROI3_collagen1647 | fibrotic | 4 | 0.68 | 1.13 |
| 40x_Fibrotic_ROI1_collagen1647 | fibrotic | 4 | 0.69 | 1.13 |
| 40x_Fibrotic_ROI2_collagen1647 | fibrotic | 4 | 0.68 | 1.13 |
| 2021-12-01-10x_fibrotic#2_collagen1.647-overview-9x5tile | fibrotic | 8890 | 2.76 | 15 |
| 2022-05-02_10X_bleomouse_LYVE1568 | fibrotic | 169 | 2.77 | 8.16 |
| 2022-05-02_10X_bleomouse-biopsy_Col1568 | fibrotic | 169 | 2.77 | 8.16 |
| 2022-05-02_10X_bleomouse_Col1568 | fibrotic | 169 | 2.77 | 8.16 |
| 10x_bleo_collagen1.568_tile2x2_V2 | fibrotic | 729 | 2.77 | 8.15 |
| bleo_collagen1568_10x_tile2x2_2 | fibrotic | 729 | 2.77 | 8.15 |
| bleo_collagen1568_10x_tile2x2 | fibrotic | 729 | 2.77 | 8.15 |
| 2022-05-17-Emphesematous-elastase-1-collagen1-568_20x_1 | emphysema | 36 | 1.38 | 2.29 |
| 2022-05-17-Emphesematous-elastase-1-collagen1-568_20x_2 | emphysema | 36 | 1.38 | 2.29 |
| 10x_emphysema1_ROI-zstack_3 | emphysema | 169 | 2.77 | 8.16 |
| 40x_emphysema1_ROI-zstack_1 | emphysema | 4 | 0.69 | 1.12 |
| 10x_emphysema1_ROI-zstack_1 | emphysema | 169 | 2.77 | 8.16 |
| 20x_emphysema1_ROI-zstack_3 | emphysema | 36 | 1.38 | 2.29 |
| 20x_emphysema1_ROI-zstack_1 | emphysema | 36 | 1.38 | 2.29 |
| 20x_emphysema1_ROI-zstack_2 | emphysema | 36 | 1.38 | 2.29 |
| 40x_emphysema1_ROI-zstack_2 | emphysema | 4 | 0.69 | 1.12 |
| 10x_emphysema1_ROI-zstack_2 | emphysema | 169 | 2.77 | 8.16 |
| 20x_emphysema1_ROI-zstack_4 | emphysema | 36 | 1.38 | 2.29 |
| 10x_emphysema1_ROI-zstack_4 | emphysema | 169 | 2.77 | 8.16 |
| 2022-02-02-20x-Emphyema#2_2 | emphysema | 36 | 1.33 | 2.29 |
| 2022-02-02-20x-Emphyema#2_4 | emphysema | 36 | 1.33 | 2.29 |
| 2022-06-10-Emphesematous-elastase-1-collagen1-568_40x_emphyyroirightsideofoverview_2x1 | emphysema | 12 | 0.69 | 1.12 |
| 2022-06-10-Emphesematous-elastase-1-collagen1-568_40x_emphyyroirightsideofoverview | emphysema | 4 | 0.69 | 1.12 |
| 10x_emphysema#1_col1_dapi_1 | emphysema | 36 | 1.31 | 2.29 |

| | | | | |
|---|---|---|---|---|
| 10x_emphysema#2_col1_overview_1 | emphysema | 10668 | 2.61 | 15 |
| 2021-12-18-10x_emphysema#1_collagen1.568-overview-7x5tile | emphysema | 6860 | 2.77 | 10 |
| 2022-06-03-Emphesematous-elastase-1-collagen1-568_20x_emphyyroi2 | emphysema | 36 | 1.38 | 2.29 |
| 2022-06-03-Emphesematous-elastase-1-collagen1-568_20x_emphyyroi | emphysema | 36 | 1.38 | 2.29 |
| 2022-06-03-Emphesematous-elastase-1-collagen1-568_40x_emphyyroi3 | emphysema | 4 | 0.69 | 1.12 |
| 2022-06-03-Emphesematous-elastase-1-collagen1-568_20x_emphyyroi3 | emphysema | 36 | 1.38 | 2.29 |
| 2022-01-26-40x-Emphyema#1_4 | emphysema | 4 | 0.69 | 1.12 |
| 2022-01-26-40x-Emphyema#1_3 | emphysema | 4 | 0.69 | 1.12 |
| 2022-01-26-10x-Emphyema#2_1 | emphysema | 169 | 2.77 | 8.16 |
| 2022-01-26-40x-Emphyema#1_1 | emphysema | 4 | 0.69 | 1.12 |
| 2022-01-26-10x-Emphyema#2_2 | emphysema | 169 | 2.77 | 8.16 |
| 2022-01-26-40x-Emphyema#1_2 | emphysema | 4 | 0.69 | 1.12 |
| 2022-05-02_10X_elastasemouse_Col1568 | emphysema | 169 | 2.77 | 8.16 |
| 2022-05-02_10X_elastasemouse-biopsy_Col1568 | emphysema | 169 | 2.77 | 8.16 |
| 2022-01-21-10x_emphysema#1_ROI_1 | emphysema | 169 | 2.77 | 8.16 |
| 2022-01-21-10x_emphysema#1_ROI_4 | emphysema | 169 | 2.77 | 8.16 |
| 2022-01-21-10x_emphysema#1_ROI_3 | emphysema | 169 | 2.77 | 8.16 |
| 2022-01-21-10x_emphysema#1_ROI_2 | emphysema | 169 | 2.77 | 8.16 |
| 2022-01-21-20x_emphysema#1_ROI_2 | emphysema | 36 | 1.38 | 2.29 |
| 2022-01-21-20x_emphysema#1_ROI_4 | emphysema | 36 | 1.38 | 2.29 |
| 2022-01-21-20x_emphysema#1_ROI_3 | emphysema | 36 | 1.38 | 2.29 |
| 2022-01-21-20x_emphysema#1_ROI_1 | emphysema | 36 | 1.38 | 2.29 |
| 40x_healthy_ROI1_collagen1647 | healthy | 4 | 0.68 | 1.13 |
| 40x_healthy_ROI2_collagen1647 | healthy | 4 | 0.68 | 1.13 |
| 40x_healthy_ROI4_collagen1647 | healthy | 4 | 0.68 | 1.13 |
| 40x_healthy_ROI3_collagen1647 | healthy | 4 | 0.68 | 1.13 |
| 2021-12-010x_healthy#2_collagen1.647-overview-9x5tile | healthy | 8890 | 2.76 | 15 |
| 2022-05-02_10X_WTmouse_LYVE1568 | healthy | 169 | 2.77 | 8.14 |
| 2022-05-02_10X_overviewtile_WTmouse_LYVE1568 | healthy | 8232 | 2.77 | 15 |
| 2022-05-02_10X_PBSmouse_LYVE1568 | healthy | 169 | 2.77 | 8.16 |
| 2022-05-02_10Xoverview_PBSmouse_Col1568 | healthy | 16129 | 2.77 | 12 |

| | | | | |
|---|---|---|---|---|
| 2022-05-02_10X_PBSmouse_Col1568 | healthy | 169 | 2.77 | 8.16 |
| 2022-05-02_10X_WTmouse-biopsy_Col1568 | healthy | 169 | 2.77 | 8.16 |

**Supplementary Table 2: Image Processing Workflow Parameters**

| Parameter Category | Parameter | Value |
|---|---|---|
| **Tiling** | Tile size | Approximately 200 µm per edge |
| **Voxel Resampling and Interpolation** | Interpolation method | Cubic interpolation (scipy.ndimage.zoom) |
| | Typical interpolation factors | 0.5, 0.5, 0.5 |
| | | 3.5, 2.0, 2.0 |
| | | 4.5, 2.0, 2.0 |
| | | 5.0, 2.0, 2.0 |
| | | 6.5, 2.0, 2.0 |
| | Final tile shape after interpolation | 92 × 92 × 92 voxels with ~200 µm per edge |
| **Semantic Segmentation** | Model | lightsheet_3D_unet_root_ds2x from PlantSeg |
| | Model training data | Light-sheet images of Arabidopsis lateral roots at 1/2 resolution |
| | Training voxel size | 0.25×0.325×0.325 µm (ZYX) |
| | Loss function | BCEDiceLoss |
| **Binarization** | Method | Otsu thresholding |

**Supplementary Table 3: 3D Image Generation Diffusion Model Parameters**

| Parameter Category | Parameter | Value |
|---|---|---|
| **Diffusion Process** | Model type | Gaussian diffusion model |
| | Timesteps | 1000 |
| | Input image size | 80 × 80 × 80 voxels (downscaled from 92 × 92 × 92 interpolated tiles) |
| | Loss function | L1 loss |
| **Augmentation** | Library | TorchIO |
| | Number of iterations | 100 |
| | Intensity rescaling | tio.RescaleIntensity(out_min_max=(0, 1)) |
| | Random elastic deformations | num_control_points=(5, 5, 5), max_displacement=(4, 4, 4), locked_borders=1 |
| | Random affine transformations | rotations: 0, 90, 180, 270 degrees independently for X, Y, Z axes |
| | Random flips | tio.RandomFlip(axes=(0, 1, 2)) |
| **Model** | Architecture | 3D unet + attention layers |
| | Base dimension (dim) | 64 |
| | Dimension multipliers (dim_mults) | (1, 2, 4, 8) |
| | Optimizer | Adam |
| | Learning rate | 1.00E-04 |
| | Training batch size | 1 |
| | Gradient accumulation | Every 2 batches |
| | Training steps | 30000 |
| | Exponential moving average (EMA) decay | 0.995 |
| | Mixed precision training | amp = True |
| | Saving frequency | 1000 steps |

**Supplementary Table 4: Mesh generation Parameters**

| Parameter Category | Parameter | Value |
|---|---|---|
| **Marching Cubes Algorithm** | Method | skimage.measure.marching_cubes |
| | Padding | 5 pixels (Probability maps and Binary Images) |
| | Binary mask dilation | scipy.ndimage.binary_dilation, 3 iterations |
| | Threshold | Isolevel, derived from the Otsu threshold |
| **Mesh Post-processing** | Library | Trimesh |
| | Steps | Removing degenerate faces (mesh.unique_faces(), mesh.nondegenerate_faces()) |
| | | General mesh processing (mesh.process()) |
| | | Fixing inverted normals (mesh.fix_normals()) |
| | | Filling any holes (mesh.fill_holes()) |

### Supplementary Figures

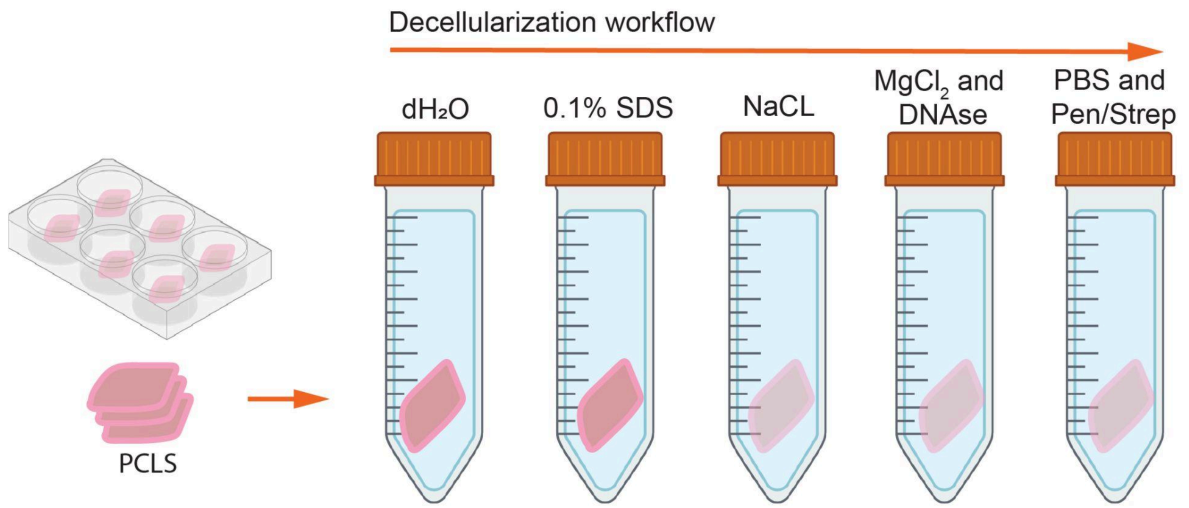


**Supplementary Fig. 1: Decellularization of mouse precision-cut lung slices (PCLS).** Schematic overview of the decellularization protocol for mouse PCLS. 300 µm thick slices were sequentially treated with distilled water (dH2O), 0.1% SDS, NaCl, MgCl2 with DNAse, and finally washed in PBS containing Penicillin/Streptomycin to remove all cellular material while preserving the extracellular matrix.

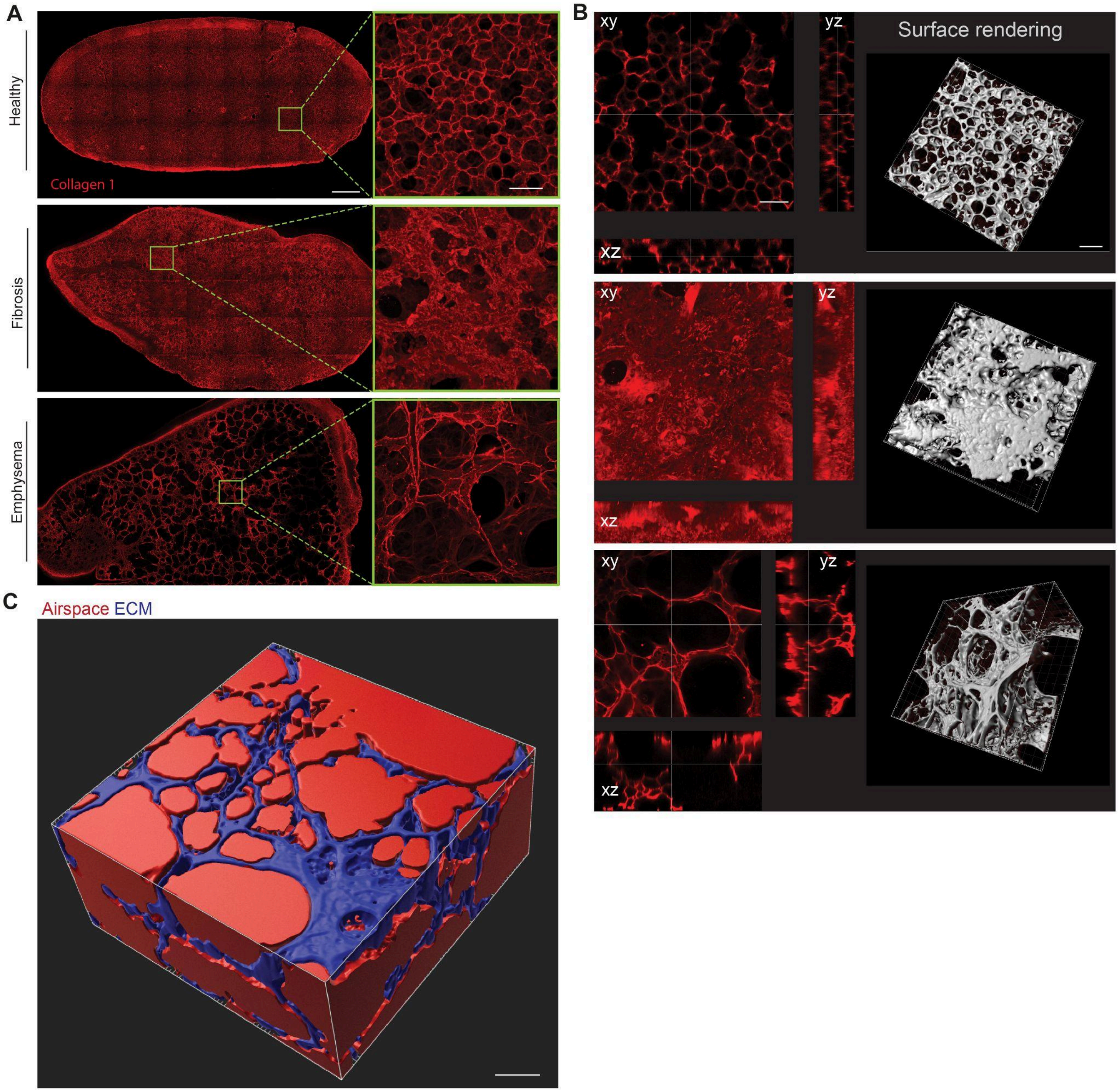


**Supplementary Fig. 2: Structural changes in decellularized precision-cut lung slices (PCLS) from healthy and diseased murine lungs. (A)** Representative tile-scanned and magnified region-of-interests (ROI) organized alveolar structure, whereas fibrotic tissue displays dense ECM accumulation and lung architecture distortion. Emphysematous slices exhibit enlarged airspaces and ECM fragmentation. Scale bar (overview) = 1000 µm. Scale bar (ROI) = 100 µm. **(B)** Orthogonal (xy, xz, yz) and 3D surface-rendered views of confocal z-stacks illustrate disease-specific ECM remodeling. Scale bars = 100 µm. **(C)** Digital segmentation of a representative mouse lung tissue volume demonstrating the process of airspace (red) vs tissue (blue) quantification for **Figure 2E**. This segmentation enabled precise volumetric quantification of disease-specific changes in the lung. Scale bar = 100 µm.

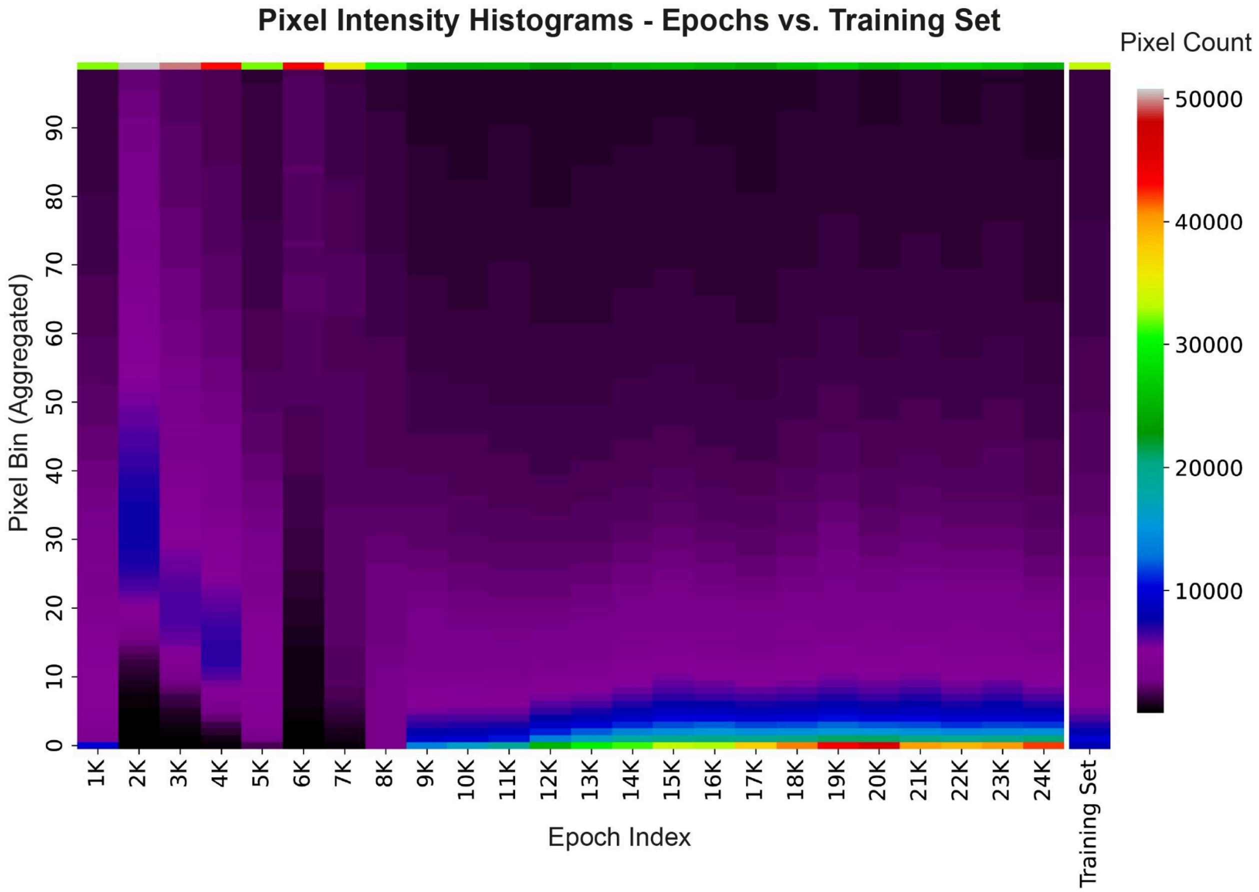


**Supplementary Fig. 3: Evolution of generated image pixel intensity distributions across training epochs** (healthy condition). The heatmap displays the frequency of pixel intensities, aggregated into 100 bins (y-axis), for images generated at different training epochs (x-axis). Each column represents the pixel intensity distribution of 100 randomly sampled images at a specific epoch (measured every 1000 epochs). The final column on the right shows the pixel intensity distribution of the original training data for direct comparison. The heatmap demonstrates a clear pattern: the images begin to show a robust separation of background and foreground around the 9,000-epoch mark, a characteristic of the real data's distribution. This separation is maintained throughout the rest of the training, although the overall quality of the distribution may not continue to improve. The data for the final epoch selection (17,000 epochs) is presented in the main text in Figure 4.

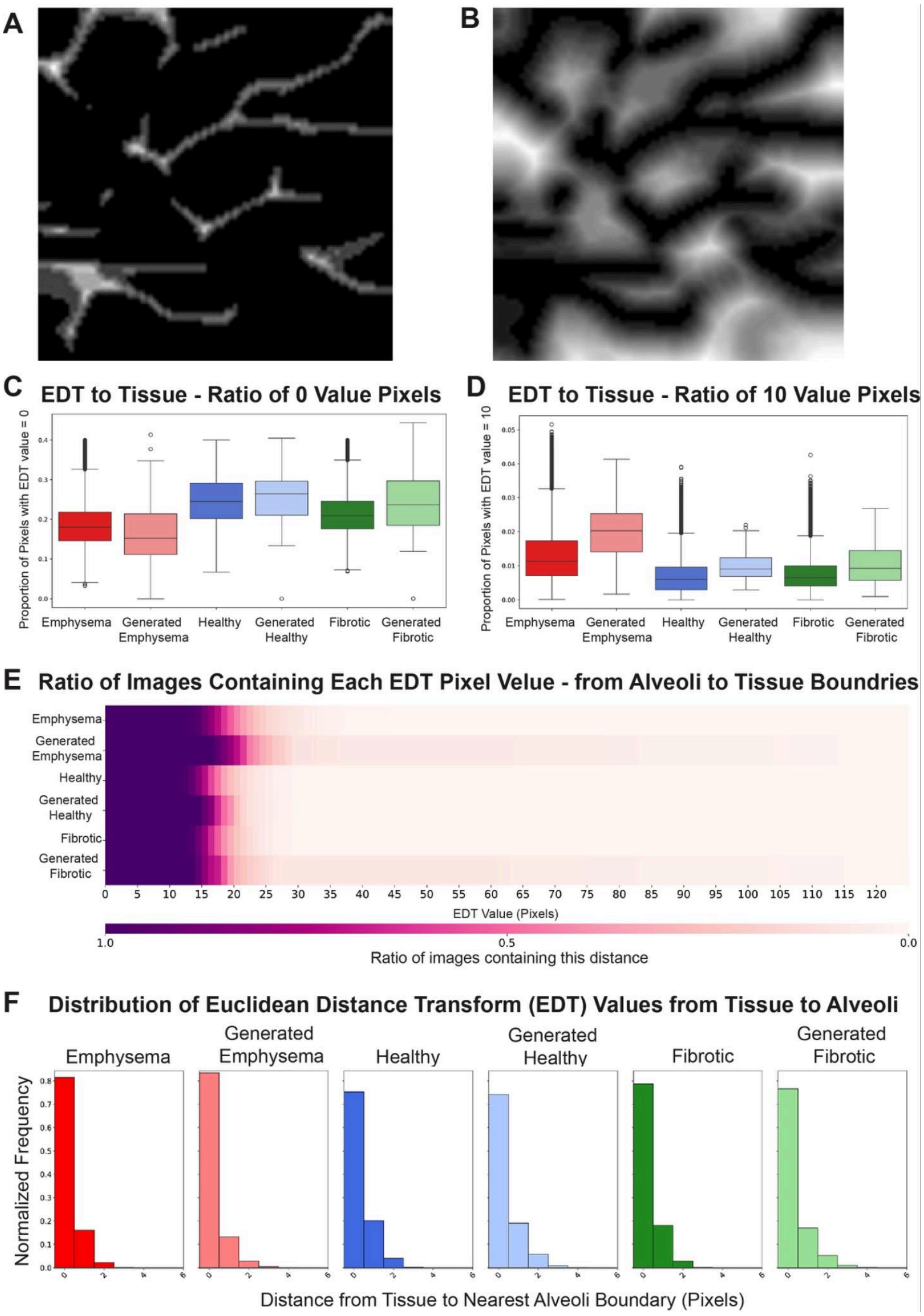
A
B
C
EDT to Tissue - Ratio of 0 Value Pixels
Proportion of Pixels with EDT value = 0
Emphysema
Generated Emphysema
Healthy
Generated Healthy
Fibrotic
Generated Fibrotic
D
EDT to Tissue - Ratio of 10 Value Pixels
Proportion of Pixels with EDT value = 10
Emphysema
Generated Emphysema
Healthy
Generated Healthy
Fibrotic
Generated Fibrotic
E
Ratio of Images Containing Each EDT Pixel Velue - from Alveoli to Tissue Boundries
Emphysema
Generated Emphysema
Healthy
Generated Healthy
Fibrotic
Generated Fibrotic
0 5 10 15 20 25 30 35 40 45 50 55 60 65 70 75 80 85 90 95 100 105 110 115 120
EDT Value (Pixels)
1.0
0.5
0.0
Ratio of images containing this distance
F
Distribution of Euclidean Distance Transform (EDT) Values from Tissue to Alveoli
Emphysema
Generated Emphysema
Healthy
Generated Healthy
Fibrotic
Generated Fibrotic
Normalized Frequency
Distance from Tissue to Nearest Alveoli Boundary (Pixels)

**Supplementary Fig. 4: Quantitative Comparison of Generated Images to Original Data Using Euclidean Distance Transform (EDT) Maps.** This figure presents a series of quantitative metrics comparing generated images to the original training data using 100 randomly sampled images per condition. All analyses utilize Euclidean Distance Transform (EDT) maps. **(A)** Example image: This 2D slice from a 3D image shows an EDT map where the distance is measured from the tissue to the alveolar boundaries. This map is generated from a binarized, segmented tissue image. **(B)** Example image: This 2D slice shows an EDT map where the distance is measured from the alveolar spaces to the tissue boundaries. This is created by flipping the binarized image and then applying the EDT. **(C)** Box plots for the ratio of zero-value pixels in the EDT to tissue. This plot compares all generated images to their corresponding original images, showing the proportion of pixels with a distance of zero. **(D)** Box plots for the ratio of pixels with a value of 10 in the EDT to tissue. This specific pixel value was chosen because it's consistently present across all EDT images and serves as a proxy for the frequency of alveoli with a size greater than this distance. The plot compares the distribution of this ratio between generated images and their corresponding original images for each condition. **(E)** Heatmap visualizing the presence of specific EDT values (alveoli to tissue) across conditions. The y-axis represents the conditions, and the x-axis represents the pixel intensity values from the EDT. The color intensity of the heatmap indicates the ratio of images per condition that contain at least one pixel with that specific EDT value. As expected, the original data from the emphysema condition shows a shift toward higher EDT values, indicating larger alveoli. However, the fibrotic images do not exhibit the expected shift toward smaller alveoli compared to the healthy tissue, which may be attributed to imaging artifacts from the challenges of imaging deep into dense tissue. For the generated images, the model consistently produces a distribution with slightly larger alveoli than the original training data across all conditions. **(F)** Normalized pixel intensity histogram. This histogram displays the normalized pixel intensity distribution of the EDT from tissue to alveoli for each condition, thus providing an estimate of tissue diameter.

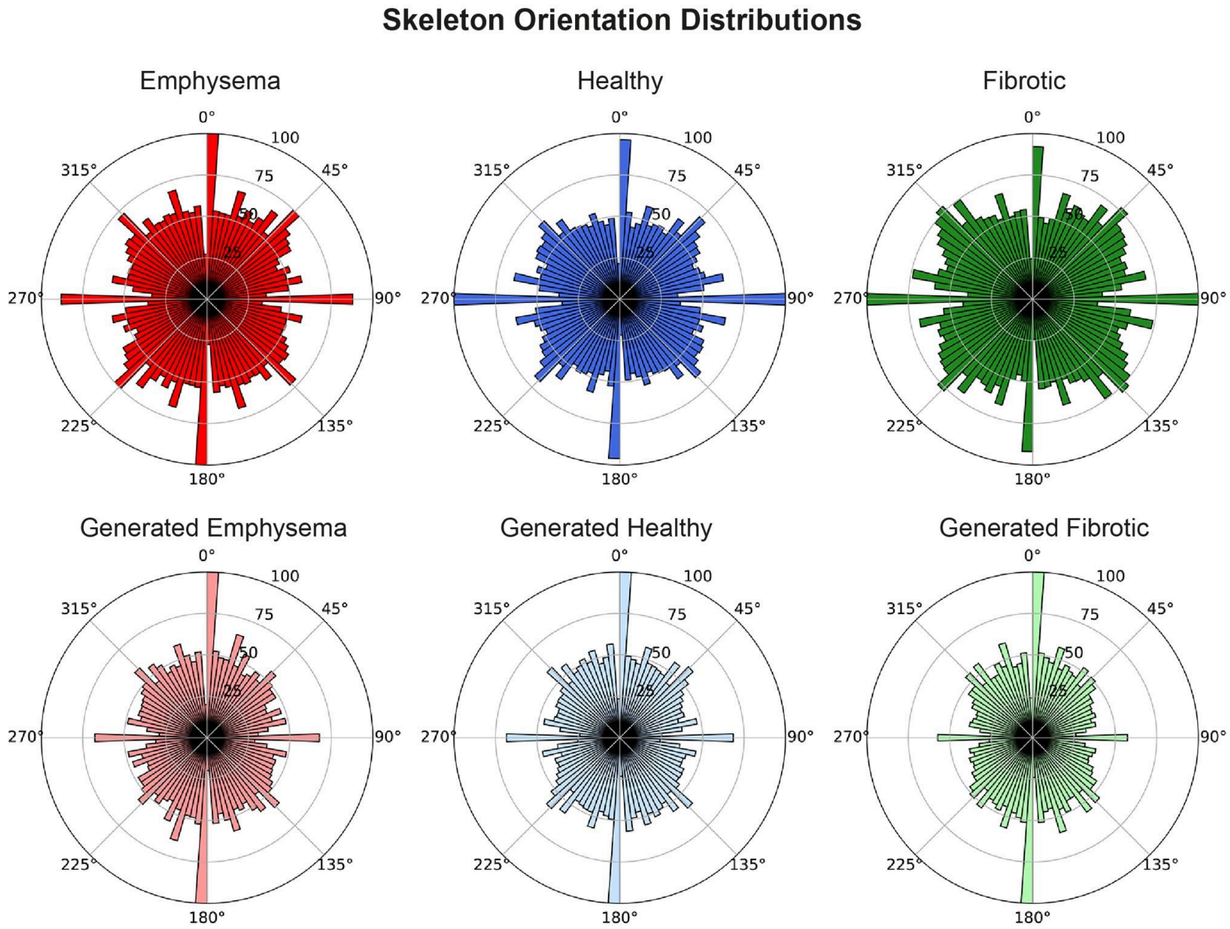


**Supplementary Fig. 5: Tissue skeleton orientation distribution.** We analyzed the orientation of filamentous structures by first creating a skeletonized representation of the binarized images. We then computed the local orientation of the skeleton at regular intervals using a sliding window approach with a 7x7 pixel window. Within each window, a 2D maximum projection was created, and Principal Component Analysis (**PCA**) was applied to the coordinates of the skeleton pixels to determine the principal direction. The orientations were filtered by a certainty threshold of 0.5 to ensure reliable measurements. The figure displays the distribution of these orientations for each condition in a rose diagram.

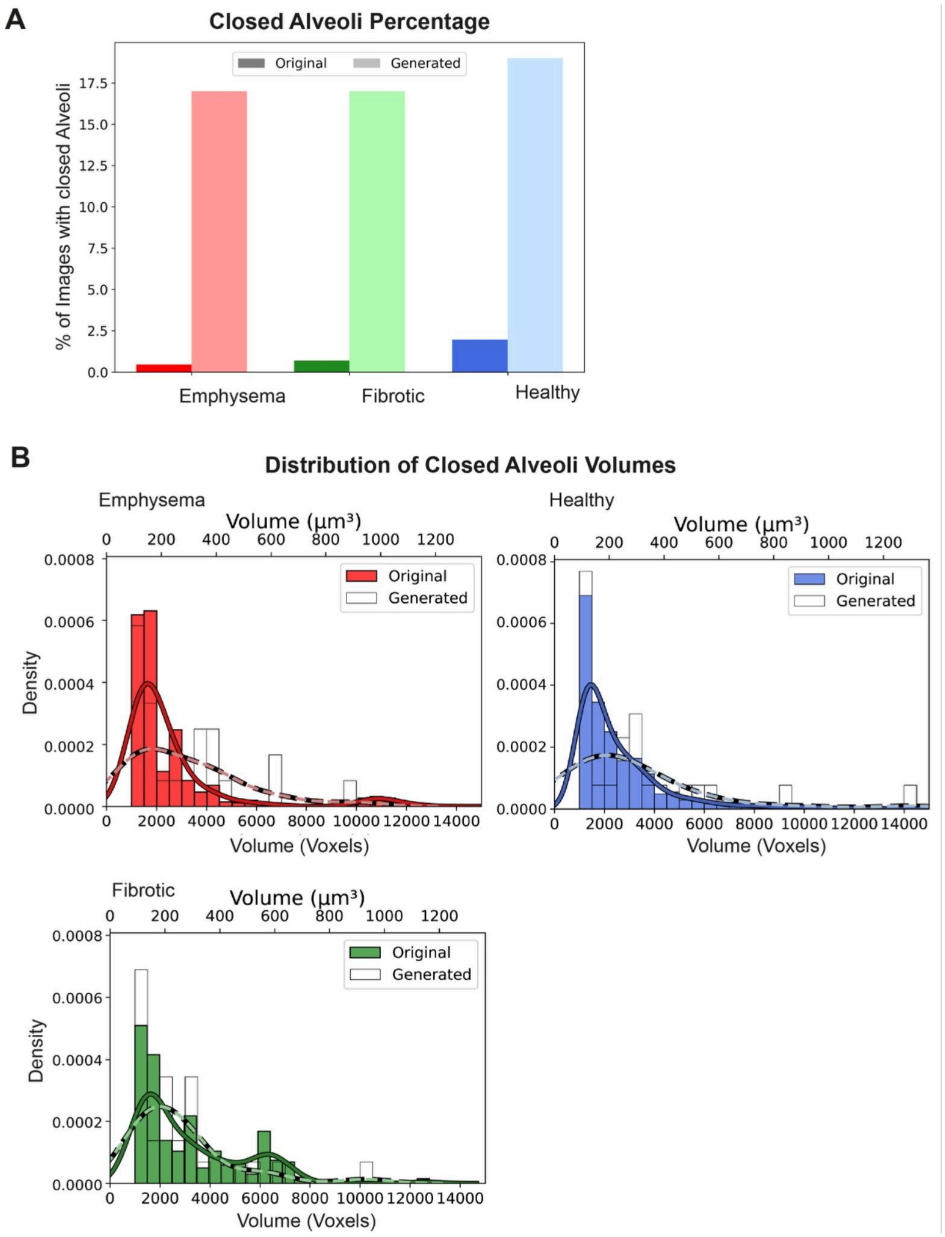


**Supplementary Fig. 6: Closed alveoli.** This figure demonstrates the prevalence and characteristics of closed alveoli (3D enclosed region), which are typically artifacts in generated models. The analysis was performed using 100 randomly sampled images per condition. **(A)** Closed alveoli percentage. This plot shows that the generated model produces a higher percentage of closed alveoli compared to the original data, a finding that is inconsistent with real tissue. **(B)** Distribution of closed alveoli volumes. To analyze the distribution of closed alveoli, we first identified and measured the volume of each closed structure. We then disregarded small enclosed volumes less than 1,000 voxels. The plot shows that after this filtering step, the volume distribution of closed alveoli in the generated images is not significantly different from the original data, suggesting the model primarily produces small artifacts. The x-axis was clipped at 15,000 voxels to better visualize the main distribution, as only 0.26% had larger volumes. Volumes are displayed in both voxels and cubic micrometers (μm³).

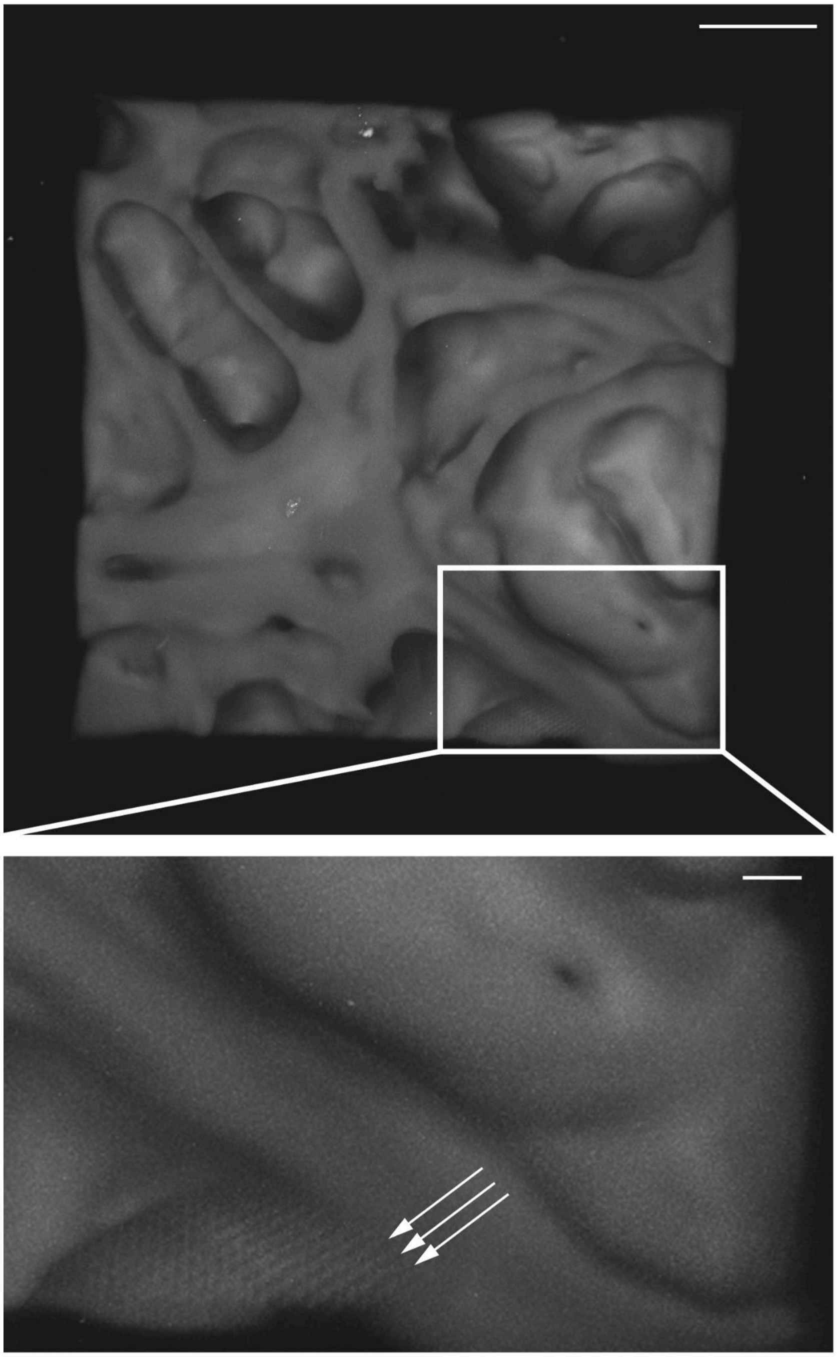

**Supplementary Fig. 7: Resolution of individual laser scan paths during two-photon polymerization of GM10 bioink.** Representative microscopy image of a 3D-printed GM10 hydrogel scaffold demonstrating micrometer precision of the printing process. **Top:** Overview of a printed lung parenchyma-like microarchitecture, illustrating overall structural similarity to real tissue. Scale bar = 50 µm. **Bottom:** Magnified view of the boxed region highlights discrete, parallel polymerized GM10-ink lines (arrows), corresponding to individual laser scan paths, which are resolved at micrometer precision. Scale bar = 10 µm.

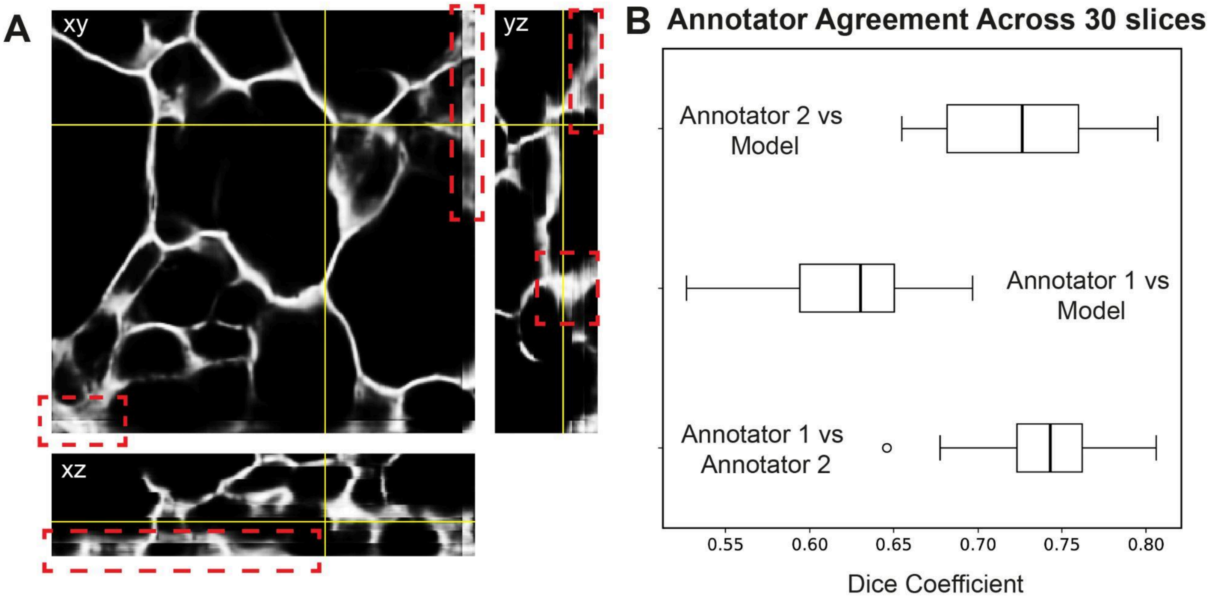


**Supplementary Fig. 8: Pretrained segmentation model selection. (A)** Artifacts from tile-based segmentation. This figure shows three orthogonal views (xy, xz, yz) of a single 3D image to illustrate artifacts from pretrained segmentation models. During the initial selection, models with a fixed tile size produced noticeable "pancake effect" artifacts along the borders of segmented regions. Examples of these artifacts are indicated by red dashed rectangles. These discontinuities occur when independently processed tiles are stitched together. Models exhibiting this issue were excluded from further consideration due to this limitation. **(B)** Quantitative model selection. To choose the final model, we compared the performance of potential candidates against human-annotated ground truth. Two independent human annotators created segmentation masks for 30 slices of a sample image. We then calculated the Dice coefficient to measure the pairwise similarity between the annotators' masks and the binarized output of each model. The selected model demonstrated a Dice coefficient comparable to the inter-annotator agreement, as shown in the pairwise comparison plot. The pairwise Dice coefficients were as follows: Annotator 1 vs Annotator 2 (mean=0.74, std=0.04), Annotator 1 vs Selected Model (mean=0.62, std=0.04), and Annotator 2 vs Selected Model (mean=0.72, std=0.04).

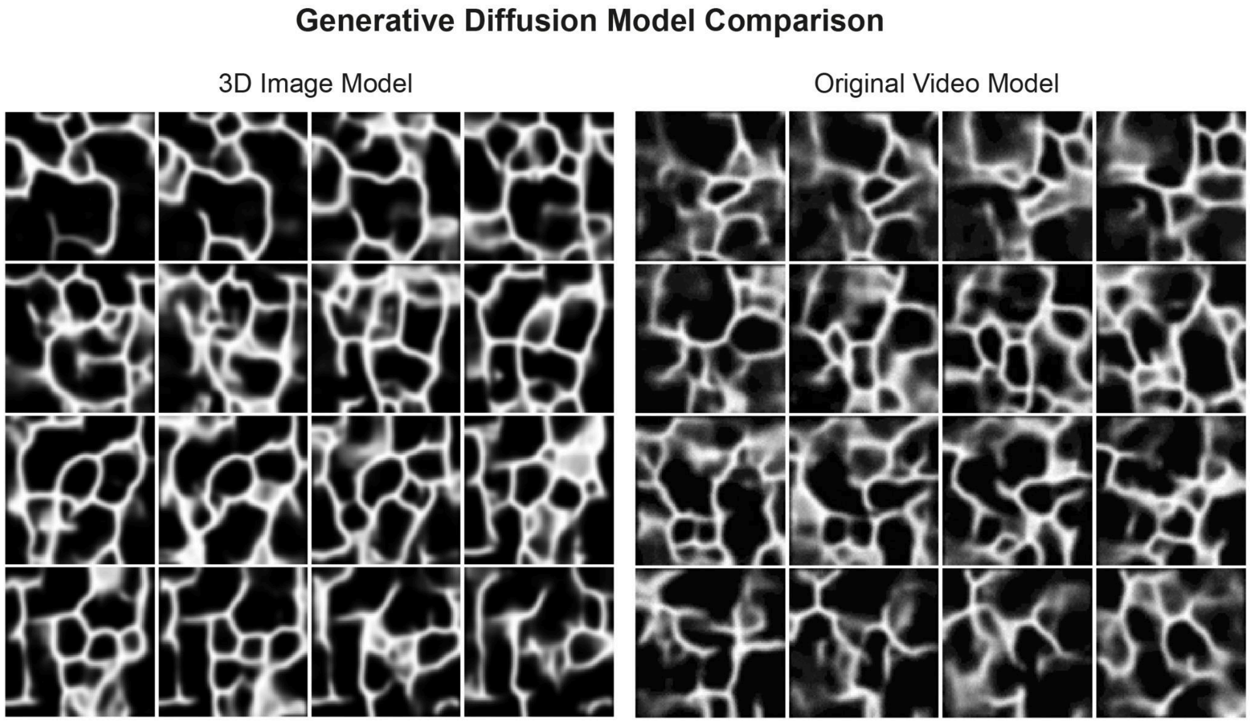


**Supplementary Fig. 9: Comparison of original video model and adapted 3D model.** To generate 3D images, we adapted an existing attention-based video generation diffusion model. The original model's architecture, consisting of a 2D spatial attention layer followed by a temporal attention layer, was modified to a single 3D attention layer to process volumetric data. Additionally, the input layer was changed to accept single-channel (grayscale) TIFF images instead of the original three-channel (RGB) GIF format. The figure displays 16 representative 2D slices (out of 92 total) from a single 3D image generated by our adapted 3D model and the original video generation model. Our adapted model produces images with greater sharpness and more well-defined borders compared to the original model, which generates blurrier output.

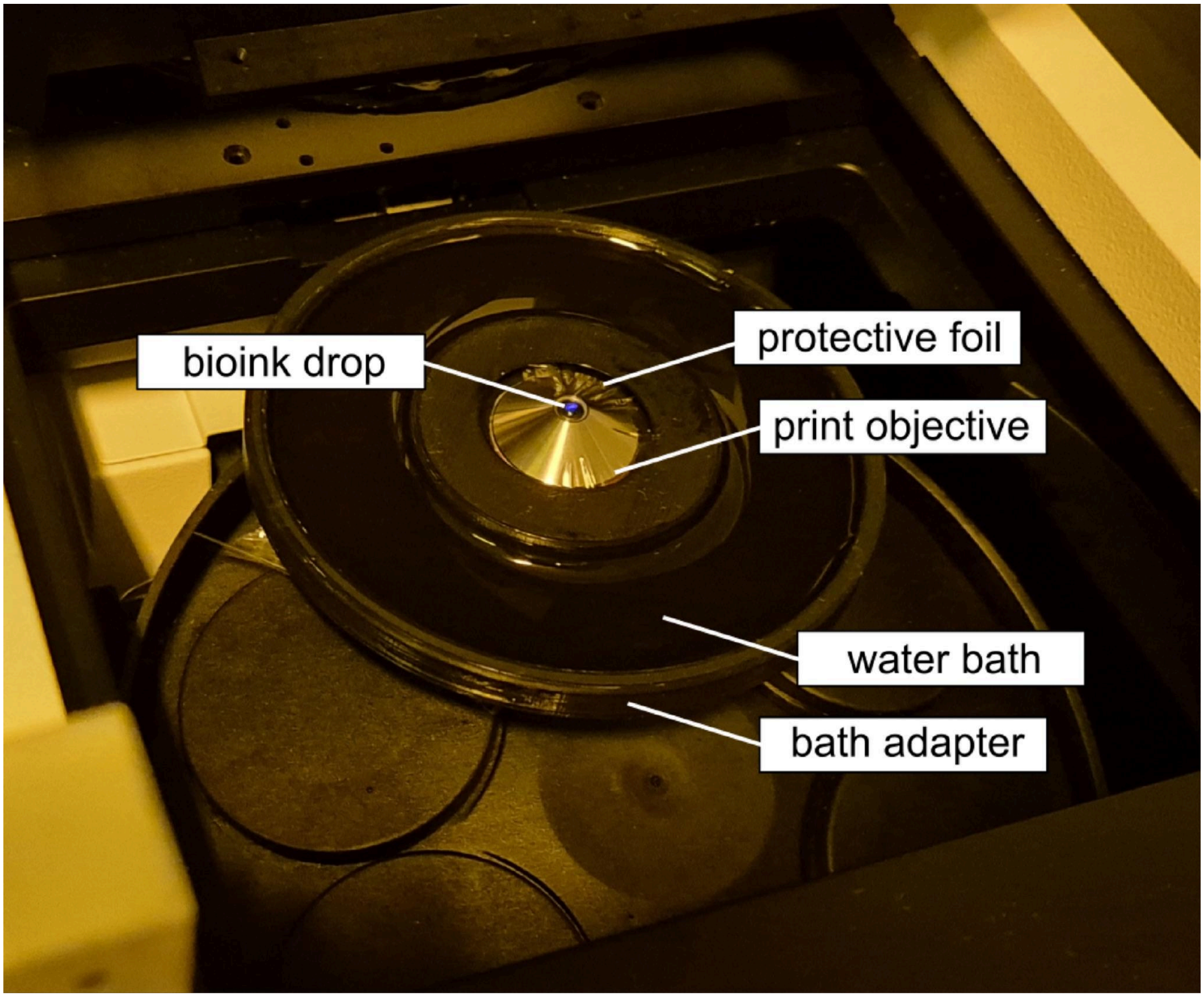


**Supplementary Fig. 10: 2-photon 3D bioprinting setup within the 3D printer.** The 25x print objective is foil-wrapped to protect the objective lens from the bioink. For fixing the foil in a stretched state, a small O-ring is attached around the objective ring. On top of the wrapped objective lens, a small drop of bioink is carefully pipetted on. Around the print objective, a custom, 3D-printed water bath adapter holds warm water to prevent the drop of the bioink from drying out.
